# Genome-scale mapping of plant transcription factor effector domains reveals conserved regulatory principles across eukaryotes

**DOI:** 10.64898/2026.10.08.757571

**Authors:** Simon Alamos, Anthony E. Sarkiss, Jacob T. Rapp, Anya Lachwani, Izaiah J. Ornelas, Maximo A. Kesselhaut, Juan D. Tibocha-Bonilla, Amy Lanctot, James K. Nuñez, Hector Garcia Martin, Patrick M. Shih

## Abstract

Eukaryotic species diverged over a billion years ago, yet our molecular view of gene regulation remains heavily skewed toward opisthokont model systems, particularly yeast and humans. Whether the protein features governing transcriptional regulation are conserved across eukaryotes or have diversified within individual supergroups remains unresolved. To provide a distinct evolutionary reference point, we used a massively parallel reporter assay in plant cells to identify transcriptional effector domains and define their sequence constraints, measuring ∼26,000 protein fragments tiling nearly all Arabidopsis transcription factors (1,859 proteins). Comparison with the activity of the same sequences in yeast revealed that activation is governed by a mixture of deeply conserved biochemical features and lineage-dependent functional differences. Likewise, plant repression domains are governed by flexible sequence grammars extending beyond conserved repression motifs. Together, these results reveal both broadly conserved and clade-specific principles governing eukaryotic transcriptional regulation.

## Introduction

Transcription factors (TFs) modulate mRNA synthesis through their activation and repression effector domains. In eukaryotes, these effector domains are often poorly conserved at the level of primary sequence and tend to be found in intrinsically disordered regions (IDRs), making it hard to determine which features governing their activity are conserved and which have diverged (*1–5*). Evaluating the same sequence in different host species is a powerful approach to reveal patterns of functional evolution independent of sequence conservation (*6–8*). However, this kind of experiment has only been conducted for a handful of transcriptional effector domains. While some activators have been shown to be functional across diverse eukaryotes, there are also many examples of effector domains that are not cross-compatible (*9–14*). Such variation in cross-species compatibility is perhaps not wholly unexpected given the substantial divergence in the basal transcriptional machinery across the eukaryotic tree (*15*, *16*). Recently, multiplexed reporter assays have used large datasets to overcome the lack of easily discernible sequence patterns in effector domains. These studies have mapped transcriptional activation and repression domains at genome scale as well as elucidated the sequence features underlying their activity (*17–24*). Yet these studies have largely sampled a single eukaryotic supergroup, Opisthokonta, leaving the sequence principles of transcriptional regulation mostly unexplored in more deeply divergent lineages. In addition, each study has probed a different set of proteins, precluding direct cross-species comparisons. This uncertainty is particularly relevant for plants, which are part of the distantly related supergroup Archaeplastida. Because of technical limitations, large-scale functional maps of plant activation domains have relied on assays in heterologous systems, particularly yeast (*4*, *9*). Consequently, we still lack a systematic view of which sequences activate transcription in plant cells in particular and which principles are conserved across eukaryotes more generally. Furthermore, despite the progress made in functionally mapping activation domains, comparable functional maps have not been generated for plant repression domains.

We recently developed ENTRAP-seq to measure the regulatory activity of thousands of protein fragments in parallel in plant cells (*25*). Here, we used ENTRAP-seq to measure the transcriptional activity of ∼26,000 40-amino-acid fragments spanning nearly all annotated TFs in the *Arabidopsis thaliana* genome (1,859 TFs). This dataset constitutes the first genome-scale functional map of TF activation domains measured in plant tissues and the first systematic functional map of plant repression domains. We compared these measurements to a prior dataset that scanned the same Arabidopsis fragments for activation in yeast (*4*). Holding thousands of protein sequences fixed while changing the cellular machinery that interprets them allowed us to decouple sequence divergence from functional conservation and study how transcriptional regulatory function has evolved across a billion years of eukaryotic divergence. We show that across the Opisthokonta-Archaeplastida divide, an activation grammar characterized by interspersed negative and aromatic residues — a stereotyped version of the canonical acidic activation domain chemistry largely described in yeast — retains activity across both hosts. However, activation domains that deviate from this chemistry can show substantial divergence in their activity in different eukaryotic hosts. In particular, we discover hundreds of activation domains that are active in plants but not in yeast. As a third reference point, many of these domains retain function in human cells, suggesting that they represent eukaryotic activation chemistries whose activity was lost or remodeled in the yeast lineage.

Our functional map of plant repressors allowed us to interrogate the role of linear sequence conservation in repression. Although canonical short linear motifs have long been thought to be key determinants of repression, we show that they are neither necessary nor sufficient for activity. Instead, repression is governed by flexible biochemical grammars characterized by blocks of aliphatic residues interspersed with charged residues within a highly basic sequence context. Thus, activation and repression are governed by distinct but flexible sequence constraints that do not rely on strong primary sequence conservation. Together, these findings show how rapid sequence turnover, deep biochemical conservation and taxon-specific evolutionary innovation can coexist in eukaryotic transcriptional effector domains.

## Results

### Genome-scale functional mapping of Arabidopsis transcriptional effector domains using ENTRAP-seq

We mapped plant effector domains at the genome scale by systematically probing all the annotated TFs of the model plant *Arabidopsis thaliana* (Fig. 1A). We followed a mapping approach based on a sufficiency criterion: a small protein fragment is considered a transcriptional effector domain if recruiting it to a reporter gene via a heterologous DNA binding domain (DBD) is sufficient to activate transcription (activation domain, AD) or repress it (repression domain, RD). We tiled the ∼2K TFs encoded in the Arabidopsis genome using 40 amino acid (aa) fragments at 20 aa intervals (Fig. 1B, Table S1). Tiles were designed to perfectly match a library previously tested in yeast cells to allow for cross-taxa comparisons (*4*). Tiles overlapping annotated DBDs were excluded to focus on regulatory regions (Fig. S1A-D). As a result, 59 TFs consisting mostly or exclusively of a DBD were excluded. This strategy yielded a library of 26,023 protein fragments from 1,859 TFs, which were split into 12 ATF sub-libraries (for *<u>A</u>rabidopsis* <u>T</u>ranscription <u>F</u>actors). Ten ATF libraries contained 2K fragments each, whereas two larger libraries contained ∼3.2K (ATF14) and 4K tiles (ATF6) to assess assay scalability (Fig. S1E). Each ATF library also contains a common set of 100 internal controls from previous datasets spanning a wide dynamic range of transcriptional activity and enabling quality assessment and cross-library harmonization (Fig. S1E).

**Figure 1.**
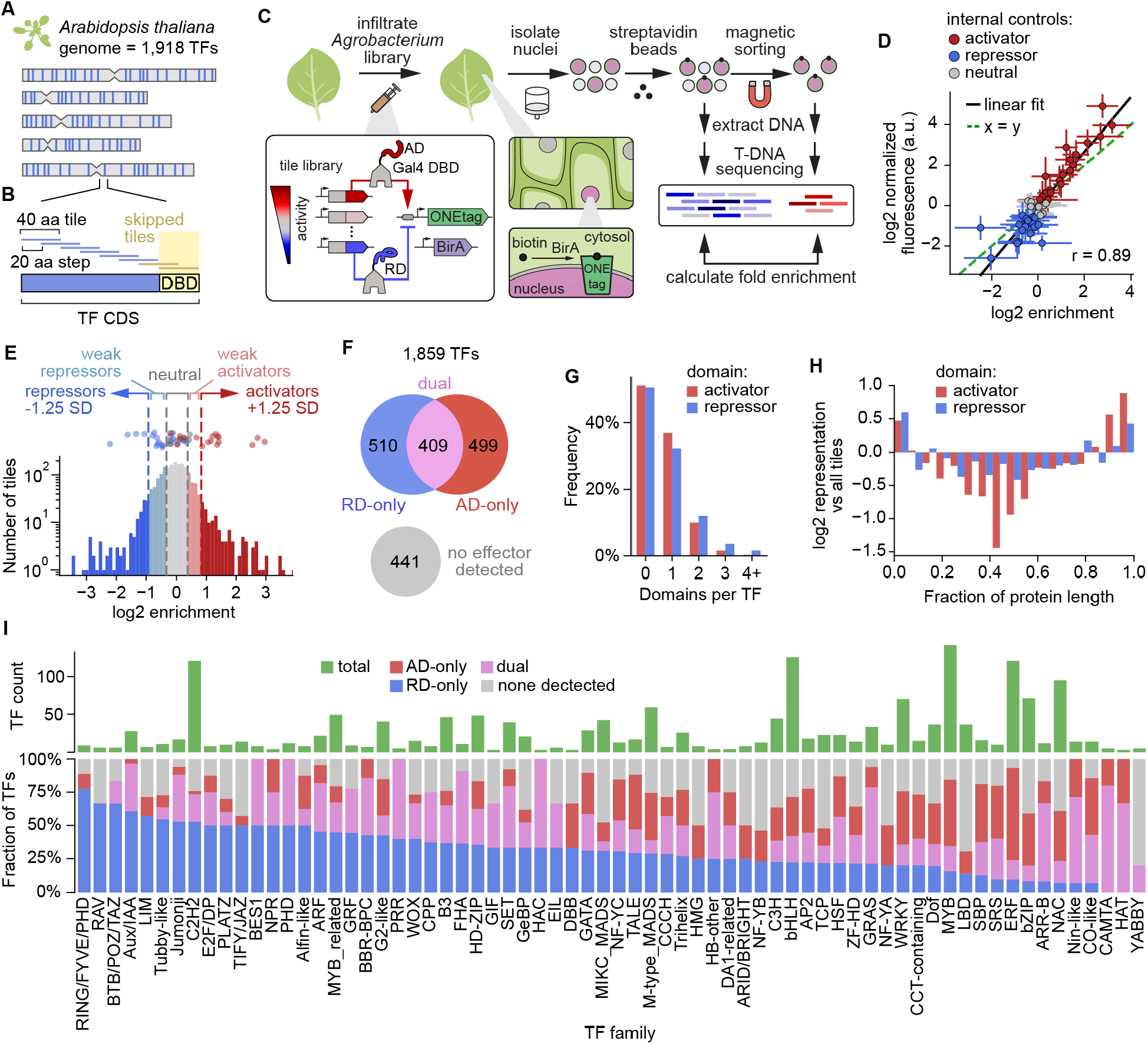
Genome-scale mapping of Arabidopsis transcriptional activation and repression domains using ENTRAP-seq. **(A)** All 1,918 annotated transcription factor (TF) genes in the *Arabidopsis thaliana* genome were selected for profiling. **(B)** TF coding sequences were tiled using 40-aa segments with a 20-aa step size. Tiles overlapping annotated DNA-binding domains (DBDs) were excluded. **(C)** ENTRAP-seq schematic. Gal4-DBD–tile fusions drive ONEtag expression in *N. benthamiana* leaves; BirA-dependent biotinylation enables magnetic sorting of nuclei, and sequencing-based pulldown/flowthrough enrichment reports tile activity, with positive values indicating activation and negative values indicating repression. **(D)** Mean ± SD transcriptional activity of 50 internal controls measured individually using a fluorescence reporter assay (y-axis) plotted against mean ± SD ENTRAP-seq enrichment of internal controls across all replicates and libraries (x-axis). Colors indicate activator, repressor, and neutral controls based on the fluorescence assay. **(E)** Distribution of enrichment values in one library (ATF1). Enrichment values of the 50 validated controls are shown on top. Dashed lines indicate enrichment cutoffs for defining qualitative activity classes. Neutral tiles fall within ±1 SD of the neutral controls. Activator and repressor tiles have enrichment values ≥1.25 SD or ≤-1.25 SD relative to the control-normalized library mean, respectively. Tiles between these thresholds were classified as weak. **(F)** 1,859 Arabidopsis TFs passing filters are classified based on the presence of repressor and/or activator tiles into RD-only, AD-only or dual (containing both AD and RD tiles). TFs in which no tiles passed classification thresholds are indicated as no effector detected. **(G)** Distribution of the number of repression and activation domains per TF. Contiguous tiles of the same category were counted as a single domain. **(H)** Distribution of the position of ADs and RDs along fractional protein length defined as amino acid position divided by protein length. **(I)** Top, in green: Number of TFs per family in the library. Bottom: Fraction of TFs per regulatory class in each family. Only families with 5 or more TFs are shown.

To measure the libraries, we used ENTRAP-seq, a multiplexed reporter assay that quantifies the transcriptional activity of thousands of protein variants in parallel directly in leaves of *N. benthamiana* (tobacco) (Fig. 1C) (*25*). Because the original implementation of ENTRAP-seq was primarily developed to profile ADs, we further optimized the assay to increase its dynamic range and simultaneously assay for both activation and repression (Fig. S2). Compared to a traditional reporter assay, optimized ENTRAP-seq accurately measured newly identified RDs of varying strengths (Fig. S2D). ENTRAP-seq biological replicates were reproducible (Fig. S3A) and an analysis of the error detected no large systematic biases in the measurement of tile enrichment (Fig. S4), allowing biological replicates to be averaged and the mean enrichment of each tile to be used as its activity estimate. To quantitatively validate the ATF libraries, we measured the activity of 50 internal control tiles using a fluorescence reporter assay and classified them as repressor, neutral, or activator relative to the Gal4 DBD alone (Fig. S1F). Their activities measured by fluorescence and ENTRAP-seq correlated well (Pearson’s *r* = 0.89, Fig. 1D), confirming that ENTRAP-seq quantitatively captures regulatory activity. The enrichment of the 100 internal controls was also reproducible between libraries (Pearson’s *r* = 0.6–0.85, Fig. S3B), supporting the integration of all libraries into a single unified dataset.

Although transcriptional activation and repression are inherently quantitative effects, many downstream analyses are facilitated by discretizing this continuum into regulatory classes based on activity thresholds. Tiles within one standard deviation of the neutral controls were classified as transcriptionally neutral, while tiles located more than 1.25 standard deviations above or below the overall mean were classified as ADs and RDs, respectively. Tiles falling between these categories were designated weak activators or weak repressors (Fig. 1E). These cutoffs were chosen to minimize AD and RD false positives (3% on average) at the expense of false negatives (35% on average), as estimated from the independently measured activity of 50 internal controls (Fig. 1E, Fig. S5). Overall, we were able to experimentally capture 97.9% of the non-control tiles. Only a small fraction of tiles were classified as RDs or ADs (approximately 6% each), consistent with the sparse distribution of effector domains observed in other systems (Fig. S5C) (*20*, *22*).

### Annotation of activation and repression domains across the Arabidopsis TF repertoire

We classified the Arabidopsis TFs into AD-only, RD-only, or dual, if they contained both an AD and an RD. Across the library, 499 (∼27%) TFs were AD-only, 510 TFs (∼27%) were RD-only, and 409 (∼22%) were dual (Fig. 1F). The high prevalence of TFs carrying both kinds of effector domains is consistent with the well-established observation that many plant TFs can activate and repress different target genes depending on context (*26*, *27*). Another 441 TFs (∼24%) lacked detectable effector domains under our assay conditions and annotation criteria. Given the estimated false negative rate, some of these neutral TFs may contain very weak effector domains that fall below our detection threshold. Alternatively, they could depend on cofactors absent from *N. benthamiana* leaves or contain larger effector domains whose activity cannot be captured in a 40-aa fragment. In general, there was good agreement between the activity of intact TF regulatory domains and their constituent tiles (Fig. S6A,B), a relationship that we further validated for 5 newly identified regulatory domains (Fig. S6C–H).

Approximately 80% of TFs containing an AD contained only one, and a similar fraction was observed for RDs (with contiguous tiles of the same sign considered a single domain) (Fig. 1G). Both ADs and RDs were strongly enriched toward the protein termini (Fig. 1H). ADs and RDs tended to occupy nearby positions within dual TFs, with 50% of opposite-function effector domains located within 80 aa of one another (Fig. S7). Mapping effector domains across TF families revealed broad differences in the occurrence of activation and repression domains (Fig. 1I). RD-only TFs are rare in large families such as ERF, MYB, WRKY, bZIP, bHLH, and NAC, whereas smaller families associated with specific developmental and signaling pathways, including Aux/IAA, JAZ/TIFY, WOX, PHD, and RAV, had mostly RD-only TFs. C2H2 TFs were a notable exception, comprising the only large family enriched in RD-containing proteins, consistent with a previous family-centric analysis (Fig. 1I) (*28*). Most families contained effector domains in the majority of their TFs, with the exception of LBD and YABBY, ∼70% of which lacked a detected AD or RD. Together, these patterns suggest a division of labor in which large TF families are dominated by activators or bifunctional regulators, while repression is concentrated in small, functionally specialized families.

Finally, we confirmed that our genome-scale functional mapping recovers previously characterized, phylogenetically conserved activator and repressor domains in the ERF, Hsf, NF-YC, DOF, and Aux/IAA families (*29–34*). Overlaying tile activity onto TF sequence-similarity dendrograms revealed widespread correspondence between evolutionary relationships and transcriptional activity across well-characterized activator and repressor clades (Fig. S8). Thus, ENTRAP-seq recovers biologically meaningful and previously validated regulatory information at genomic scale.

### A genome-scale cross-host experiment reveals conservation and divergence patterns

Large-scale functional studies have identified sequence features that predict transcriptional activation in individual eukaryotic species (*4*, *17–23*). However, these relationships have been learned within individual experimental systems, each using a different protein library. It thus remains unclear which features reflect general properties of eukaryotic activation and which depend on lineage-specific cellular machinery. We addressed this limitation through a genome-scale cross-host experiment, combining our in-planta measurements with a previous dataset in which the same Arabidopsis tiles were assayed for transcriptional activation in *S. cerevisiae* (Fig. 2A) (*4*). Plants and fungi belong to distinct eukaryotic clades (i.e., Archaeplastida and Opisthokonta), which diverged at least one billion years ago (*35*). Thus, for thousands of identical protein sequences, we could directly ask which features retain function across a billion years of eukaryotic divergence and which depend on lineage-specific cellular machinery (Fig. 2A).

**Figure 2.**
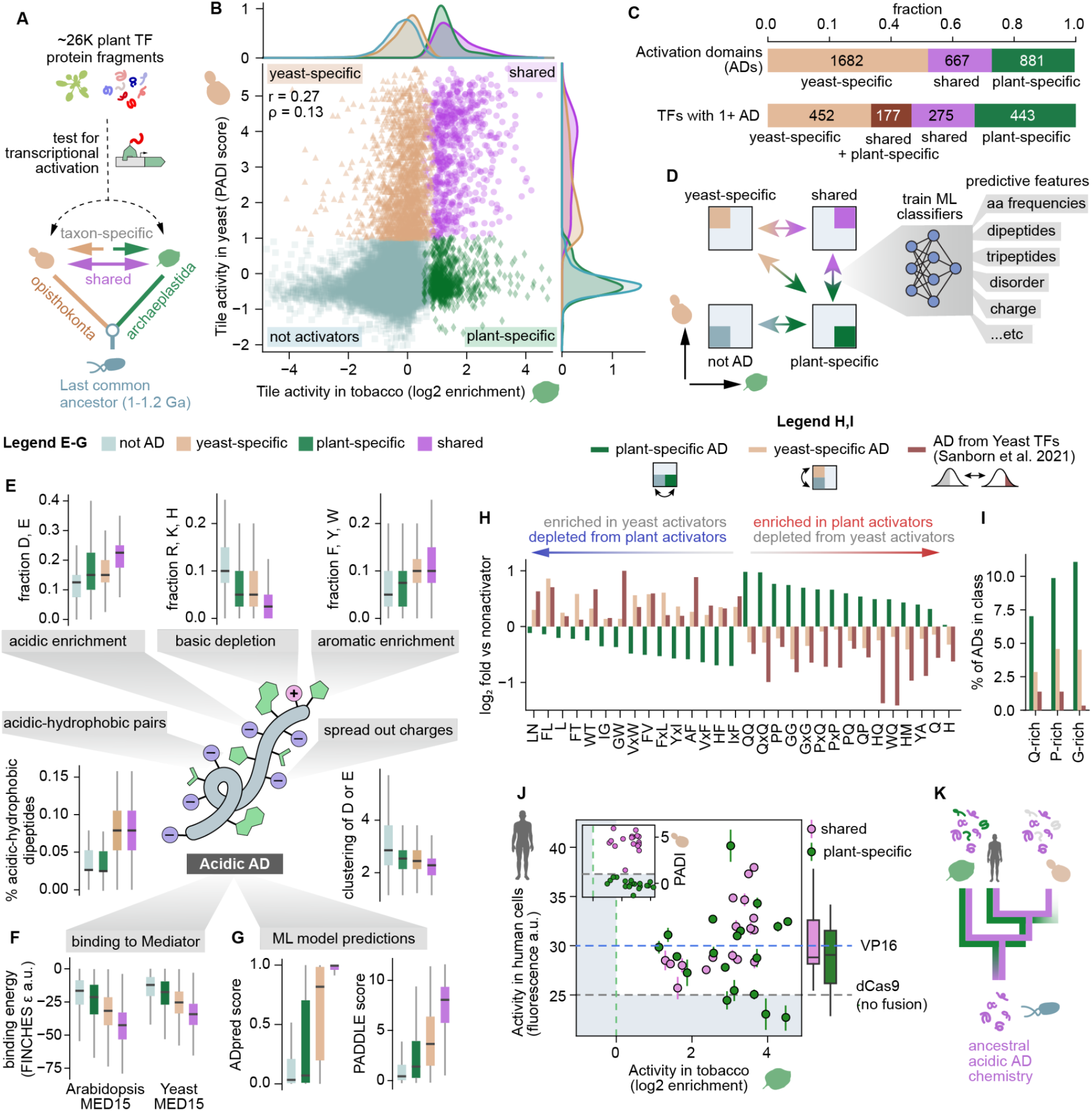
Elucidating patterns of conservation and divergence across eukaryotic activation domains. **(A)** Schematic of the experiment. 40-aa protein tiles from Arabidopsis TFs are tested in tobacco (this study) or yeast (*4*), which shared a common ancestor approximately 1 billion years ago. **(B)** Transcriptional activity of the Arabidopsis TF tile library in yeast (y-axis, PADI scores) and tobacco (x-axis, ENTRAP-seq). In yeast, tiles with a PADI score ≥ 1 were previously classified as ADs. Classifying tiles as ADs or not ADs in each experiment defines 4 quadrants, resulting in 3 AD classes: yeast-specific, plant-specific, and shared (ADs in both). Non-AD tiles are repressor, weak repressor or neutral in tobacco and with a PADI score <1. Tiles in the weak activator category in tobacco were not included because their assignment to these quadrants is ambiguous. Histograms show the density distribution of values for each quadrant along each axis. **(C)** Top: fraction of ADs that belong in each activation class in (B). Bottom: TFs with at least one tile in at least one of the AD quadrants were classified according to the class of the AD contained in the TF. In brown (shared + plant-specific) are TFs with no yeast-specific tiles that contain at least one shared AD and at least one plant-specific AD. **(D)** Machine learning approach to determine the features distinguishing classes. Pairs of AD classes were classified from one another using random forests trained on interpretable sequence features. Plant-specific ADs were classified from non-ADs using regressors trained on the same features. **(E)** Features associated with the acidic AD biochemical grammar identified in (D) are shown for the AD classes in (B). **(F)** Predicted binding energy between ADs in each class in (B) and Arabidopsis and yeast Mediator subunit 15. **(G)** AD prediction scores from two models trained in yeast to predict acidic ADs. **(H)** Fold change in the abundance of single amino acids, dipeptides, and dipeptides with a gap (‘x’) with respect to their abundance in non-activators (*20*). Shown are peptides that were simultaneously enriched in plant-specific ADs and depleted from yeast-active ADs, or vice versa. **(I)** Percentage of ADs belonging to each class (defined as at least 15% of residues being indicated aa). **(J)** Mean ± SD transcriptional regulatory activity of tiles from Arabidopsis TFs in human cells (y-axis) and tobacco (x-axis). In human cells, tiles were fused to dCas9 and used to activate the endogenous EPCAM gene, using 2 biological replicates and VP16 as a positive control. Activity was measured using flow cytometry of cells stained with labelled EPCAM antibodies. **(K)** Proposed model for the evolution of plant-specific and shared ADs. Box plots in (E)-(G) and (J) show the median ± IQR, whiskers correspond to minimum and maximum values excluding outliers.

Comparing the activity of TF fragments in yeast (PADI scores (*4*)) and tobacco (ENTRAP-seq enrichment) revealed a moderate overall correlation (Fig. 2B and Fig. S9A, Spearman ⍴ = 0.13, Pearson r = 0.27) and the odds that a fragment activated in yeast increased with its activity in plants (Fig. S9B). Nevertheless, the overlap between activators in the two systems was only partial. Among yeast activators (defined as PADI ≥ 1), only about one quarter were also classified as plant activators (Fig. 2C). Conversely, approximately half of plant activators failed to activate in yeast (PADI < 1, Fig. 2C). A similar lack of agreement was found at the TF level, with ∼900 genes carrying ADs found only in one model organism and not the other (Fig. 2C). These discrepancies were robust to activity thresholds and could not be readily ascribed to differences in disorder, synthetic TF design, or cell type dependence (Fig. S9C–F). Limited cross-compatibility was not specific to fragments of plant origin. About half of 25 strong ADs from the yeast genome were not activators in plants (Fig. S10). We therefore defined three operational classes: 1) shared ADs, active in both hosts; 2) plant-specific ADs, active only in tobacco; and 3) yeast-specific ADs, active only in yeast. These definitions describe activity in the two distinct host assays and do not imply activity or inactivity in other eukaryotes.

### Conserved and lineage-specific activation protein features

To identify the specific features that distinguish AD classes, we trained random forest classifiers on interpretable sequence features such as biophysical parameters, amino acid frequencies, and counts of dipeptides (Fig. 2D, Table S2). To find features specifically associated with activation along the plant axis, we trained a regression model on the same feature set using only tiles that do not activate in yeast (i.e., keeping only tiles with PADI score <1). We complemented this plant-specific regressor with a regressor trained on all plant data, regardless of yeast scores. Models performed well on held-out data (Table S3, Fig. S11-S13) and relied mostly –although not exclusively– on features characteristic of acidic ADs, a well-established and widespread type of eukaryotic activation domain (*2*, *3*, *17–23*, *36*, *37*). These features included enrichment of negatively charged and aromatic residues, depletion of positive charge, and patterns involving negative and hydrophobic residues (Fig. 2E, Fig. S11-S13). Across these features, shared ADs showed the strongest bias, yeast-specific ADs were intermediate, and plant-specific ADs showed the weakest signal (Fig. 2E, Fig. S11). Thus, the three AD classes are organized along a common axis corresponding to increasing conformity to canonical acidic AD chemistry. Mechanistically, acidic ADs are thought to act in part through binding to Mediator subunit 15 (MED15) (*20*, *38*, *39*); we therefore used FINCHES to predict binding of each AD class to Arabidopsis and yeast MED15 (*40*). Predicted binding recapitulated the same progression, with plant-specific ADs predicted to bind very weakly, yeast-specific ADs being intermediate, and shared ADs binding strongly (Fig. 2F, Fig. S9G). Previous machine learning models trained on yeast data to predict acidic ADs also placed the three classes in the same order (Fig. 2G, Fig. S9H) (*4*, *19*, *20*, *24*). Thus, sequence features, predicted MED15 binding, and independent models of AD activity all place the three classes along the same canonical acidic AD axis.

The ordering from neutral to plant-specific, yeast-specific, and shared ADs along the acidic AD axis may suggest that host specificity simply reflects differences in sensitivity to this chemistry. However, yeast-specific ADs are more canonically acidic than plant-specific ADs yet activate only in yeast, suggesting that additional sequence determinants contribute to host specificity. We therefore searched for features depleted from plant-specific ADs but enriched in both yeast-specific and endogenous yeast ADs. Peptides containing F and the aliphatic residues V, I, and L showed this pattern (Fig. 2H). This was largely consistent with the negative contribution of these features in our regression models for plant activation (Fig. S12, Fig. S13), even though F and L are known to contribute to yeast activation (*41*). Conversely, peptides containing Q, G, P, and H were enriched in plant-specific ADs but depleted from tiles that activate in yeast, be it from plant or yeast origin (Fig. 2H). Notably, Q-, G-, and P-rich ADs are common in metazoans but rare and generally inactive in yeast (*2*, *13*, *20*, *42*). Notably, our plant-based assays identified the well-known Q-rich ADs of activator ARFs, which were inactive in yeast (Fig. S14) (*4*, *43*). Each of these classes represented ∼10% of plant-specific ADs but <2% of endogenous yeast ADs (Fig. 2I). This raised the possibility that many plant-specific ADs are actually broadly functional eukaryotic ADs that yeast fails to recognize. To test this idea, we measured the transcriptional activity of 34 strong and randomly selected shared and plant-specific ADs in human cells. In contrast to their behavior in yeast, these tiles exhibited similar activities in human cells, with many plant-specific ADs activating at least as strongly as VP16 (Fig. 2J).

Together, these results support a model in which an ancestral and conserved acidic AD chemistry evolved in a lineage-specific manner, with partial loss or remodeling in *S. cerevisiae* (Fig. 2K). Because the acidic AD grammar captured by shared ADs is present in plant, fungal, and animal activation domains, the most parsimonious explanation is that this AD chemistry predates the divergence of these lineages, although recurrent convergence cannot be excluded. Recent cross-kingdom profiling of chromatin regulators similarly found greater functional concordance between plant and human cells than between either system and yeast (*44*). Regarding yeast-specific ADs, their acidic AD chemistry may be counteracted by features that are interpreted as inhibitory in plants. In particular, while L contributes to the hydrophobicity of acidic ADs in yeast (*41*), in plants it appears to have an inhibitory effect (Fig. S12, Fig. S13). The recurring enrichment of aliphatic residues, particularly V, I and L, raises the possibility that these residues contribute to a previously unrecognized repression-biasing grammar in plants.

### Functional repression motifs emerge from a distributed sequence grammar

Our cross-host analysis of activation domains showed that regulatory function can be deeply conserved despite weak primary-sequence conservation, through preservation of a distributed biochemical grammar. Repression provides a complementary intra-species test of this relationship between sequence conservation and function. Unlike activation, the prevailing view of TF-mediated repression in eukaryotes remains strongly centered on linear sequence conservation. In this framework, direct repression by TFs is mediated by conserved short linear motifs of 4-10 aa (sometimes referred to as SLiMs) that recruit co-repressors (*18*, *32*, *45–48*). In plants, the best-known such motifs are the ERF-associated amphiphilic repression (EAR) motifs, which recruit TOPLESS/TOPLESS-RELATED (TPL/TPR) co-repressors, although several less prevalent repression-associated motifs have also been characterized (Table S4) (*32*, *48–52*). These motifs are therefore considered major determinants of repression: mutating their conserved residues can abolish activity, and fusion of repression motifs can convert activators into repressors (*32*, *53*). In plants, however, this motif-centric picture has not been systematically examined using repression assays, and we lack a comprehensive sequence-to-function framework for plant repression domains (*49*, *54*, *55*).

Consistent with the motif-centric view, tiles containing at least one canonical motif have more negative enrichment values than tiles lacking known motifs (Fig. 3A, Fig. S15A,B) and motifs are enriched among repressors, both at the level of tiles and TFs (Fig. 3B, Fig. S15A,B). However, about 80% of RDs lacked canonical motifs, and ∼75% of TFs containing at least one RD do not have canonical motifs in any of their RDs (Fig. 3B, Fig. S15A), suggesting that our understanding of plant RDs is largely incomplete. Interestingly, many motif-containing tiles showed no detectable repressor activity (Fig. 3A, Fig. S15A). For example, ∼70% (491/690) tiles containing the LxLxL EAR motif were not repressors. Hence, in our assay, the presence of a canonical motif is neither necessary nor sufficient for repression, mirroring findings in fruit flies (*18*). This incomplete association could not be explained by predicted structural disorder or by motif position along the tile (Fig. S15C,D). The effect size of canonical motifs on ENTRAP-seq enrichment ranged from modest (Cliff’s δ = -0.26 for the TIFY motif) to extremely strong (Cliff’s δ = -0.97 for the TLxLFP motif, where x = any amino acid) (Fig. 3C).

**Figure 3.**
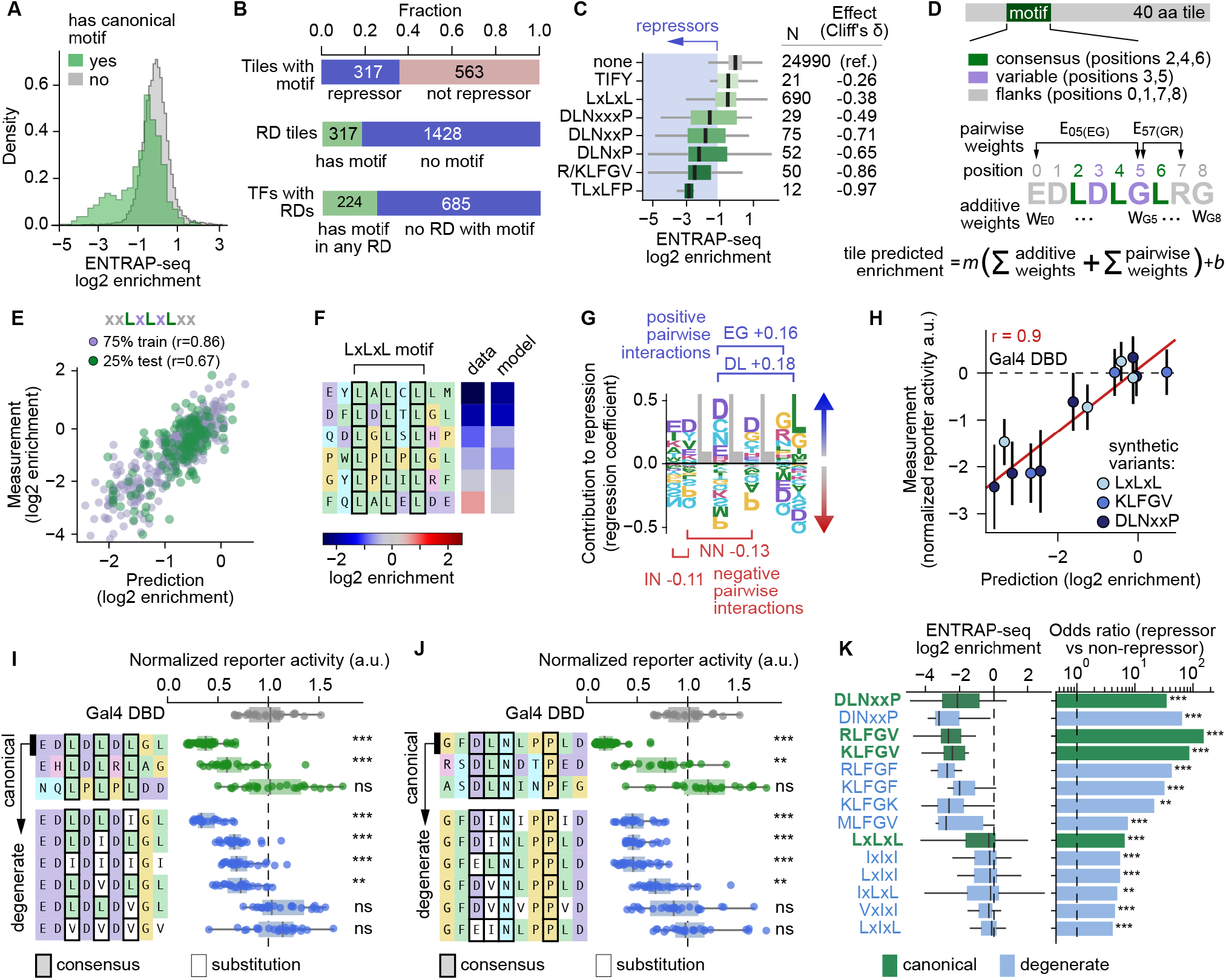
Systematic dissection of repression motifs as flexible and quantitative units. **(A)** Density histograms showing the distribution of ENTRAP-seq enrichment values for tiles with a canonical repression motif (green), or without (gray). **(B)** Top: Tiles that contain at least one canonical repression motif partitioned into repressors, or non-repressors. Middle: Fraction of repressor tiles that do or do not have a canonical repression motif. Bottom: Repressor TFs (i.e., TFs with at least one RD) partitioned between those which have a canonical motif in their RD and those which do not. **(C)** Boxplots showing the ENTRAP-seq enrichment values of tiles containing each canonical motif. **(D)** Ridge-regression model in which amino-acid identities at variable motif and ±2 flanking positions contribute additive and pairwise terms that predict tile enrichment. Weights are shared across all occurrences and the sum of all weights determines the ENTRAP-seq enrichment of the tile carrying a specific motif. **(E)** Measurements (y-axis) versus predictions (x-axis) for the model in (D) applied to the EAR motif LxLxL. **(F)** Examples of motifs from Arabidopsis tiles displaying measured and predicted differences in repression strength. Consensus positions are outlined. **(G)** Repression weights logo for LxLxL with two flanks. The height of each amino acid corresponds to the magnitude of its linear weight either contributing to repression (positive weights), or against it (negative weights). The two strongest pairwise interactions favoring or disfavoring repression are also shown. **(H)** Mean ± SD measured versus predicted repression strength of synthetic motifs tested as Gal4 DBD fusions in a fluorescence assay. **(I, J)** Fluorescence reporter assay of Gal4 DBD fusion peptides encoding EAR motifs not found in the library, (I) LxLxL and (J) DLNxxP. On top, peptides with canonical motifs and different repression strengths (green). The strongest canonical peptide was mutated to generate the degenerate peptides shown in blue at the bottom. Degenerate peptides carry substitutions that disrupt the canonical motif. **(K)** Association between degenerate motifs and repression. Left: box plot of ENTRAP-seq enrichment values for each motif. Right: odds ratio between belonging to repressors or non-repressors for each motif. Box plots in (C) and (I)-(K) show the median ± IQR, whiskers correspond to minimum and maximum values excluding outliers. In (I) and (J), *** p<0.001, ** p< 0.01, * p<0.05, ns not significantly lower using a one-sided Welch’s t-test against Gal4-DBD control. In (K), *** p<0.001, ** p< 0.01 Fisher’s exact test for enrichment in repressors versus neutral, weak activator and activator, combined. In (H)-(J) N = 24 (4 plants with 6 leaf discs per plant).

Since canonical motifs are not binary encoders of repression, we asked which additional sequence features underlie quantitative differences in strength. We hypothesized that these differences are encoded by the nonconsensus positions within the motif, their pairwise interactions, and the flanking sequence (*50*, *56*). To test these ideas, we devised a simple linear model for the LxLxL motif with two flanking positions on each side (i.e. xxLxLxLxx), in which each x position contributes an additive activity weight determined by its amino acid identity, while each pair of residues contributes a pairwise weight (Fig. 3D). Weights act as constants shared across the whole library and the sum of all weights corresponds to the repression strength of the tile containing a given motif variant (Fig. 3D). We trained this model on the 690 tiles containing the LxLxL EAR motif and found good predictive performance on held-out data (Pearson *r* = 0.67) (Fig. 3E, F). Visualization of the fitted additive coefficients as a repression weight matrix logo revealed strong amino-acid preferences at the internal free ‘x’ positions and also at the flanking residues (Fig. 3G). Consistent with previous observations, negative interspersed residues and a fourth L strongly promote repression (Fig. 3G) (*50*, *53*). We obtained similar predictive power for the DLNxxP EAR motif and the KLFG motif using analogous models despite the substantially smaller sample sizes (Fig. S15E,H). The model also accurately predicted the activity of synthetic sequences absent from the training dataset (r = 0.9) (Fig. 3H). Across motifs, the weights of some pairwise interactions were comparable in magnitude to those associated with the core positions (Fig. 3G; Fig. S15G,I). Consistent with this, removing flanking residues or pairwise interactions from the model substantially reduced predictive performance for LxLxL (Fig. S15J,K), suggesting that both local sequence context and residue-by-residue interactions contribute to repression strength.

Given the contribution of variable and flanking residues to motif strength, we asked whether an optimal sequence context could functionally compensate for mutations in consensus motif positions. If so, degenerate motifs with optimal contexts should outperform consensus motifs with detrimental contexts. We took the most potent of our synthetic EAR variants and systematically mutated the anchor motif residues to chemically similar amino acids. Many of these degenerate peptides retained activity and many were stronger than synthetic peptides containing canonical motifs (Fig. 3I, J). We obtained comparable results with mutants of SRDX, a well-known synthetic optimized EAR peptide, and with the KLFG motif (Fig. S16) (*57*). Consistent with these results, across the whole library, degenerate versions of several motifs are associated with repression at least as much as their canonical counterparts (Fig. 3K).

Thus, just as the sequence context modulates the activity of a motif, it can also relax the sequence requirements thought to define repression motifs in the first place. Together, the importance of variable and flanking residues, the tolerance of substitutions of consensus residues, and the absence of identifiable motifs in many repression domains point to repression being encoded by a far more flexible and distributed sequence grammar than a strict motif-centric view would suggest. This view is reminiscent of recent findings showing the flexible and context-dependent nature of motifs in yeast and animal ADs (*5*, *58*).

### A newly described repression motif

Because about 80% of RDs lacked canonical motifs, we hypothesized there may be novel motifs that have not yet been described. To test this, we searched for 4-mers enriched among repressor tiles after removing all tiles containing canonical motifs (Fig. 4A). The most significantly enriched motifs, including FDLN and LFGK, resembled degenerate versions of the DLN-type EAR and R/KLFGV canonical repression motifs, consistent with our finding that permissive sequence contexts can relax strict motif requirements. Nonetheless, these enriched partial motifs are present in only 71 out of 1,428 RDs lacking canonical motifs. The analysis also identified a small number of new candidate motifs (Fig. 4A). The three strongest candidates, HARF, ARFR and RFRR, overlapped within the same RDs, suggesting that they represent a single underlying motif. We therefore tested the corresponding 6-aa peptide, HARFRR. Among all new motif candidates, only HARFRR functioned as an autonomous repression motif, with a strength comparable to the potent synthetic SRDX peptide (Fig. 4B). Furthermore, fusion of HARFRR to the activator ARR12 completely abolished activation of its native target promoter in a transient assay (Fig. 4C) (*59*). The HARFRR sequence is conserved in N-terminal RD tiles of five group IId WRKY TFs which are known negative regulators containing an N-terminal repression domain (Fig. 4D) (*60*, *61*). Mutation of the motif (FRR→ADE) significantly reduced repression by the N-terminal regulatory region of WRKY7, although it did not completely abolish it (Fig. 4E). Together, these results establish this peptide as a repression motif. Notably, the HARFRR motif accounted for only 6 out of 1,428 motifless repression domains. Thus, the vast majority of plant RDs still cannot be readily explained by traditional conserved motifs of four or more residues. Although additional motifs restricted to one or a few TFs almost certainly exist, they are unlikely to account for most motifless repression domains (*62*). Instead, these results support a model in which repression, like activation, can be encoded by sequence constraints that are hard to detect using approaches based on linear sequence conservation.

**Figure 4.**
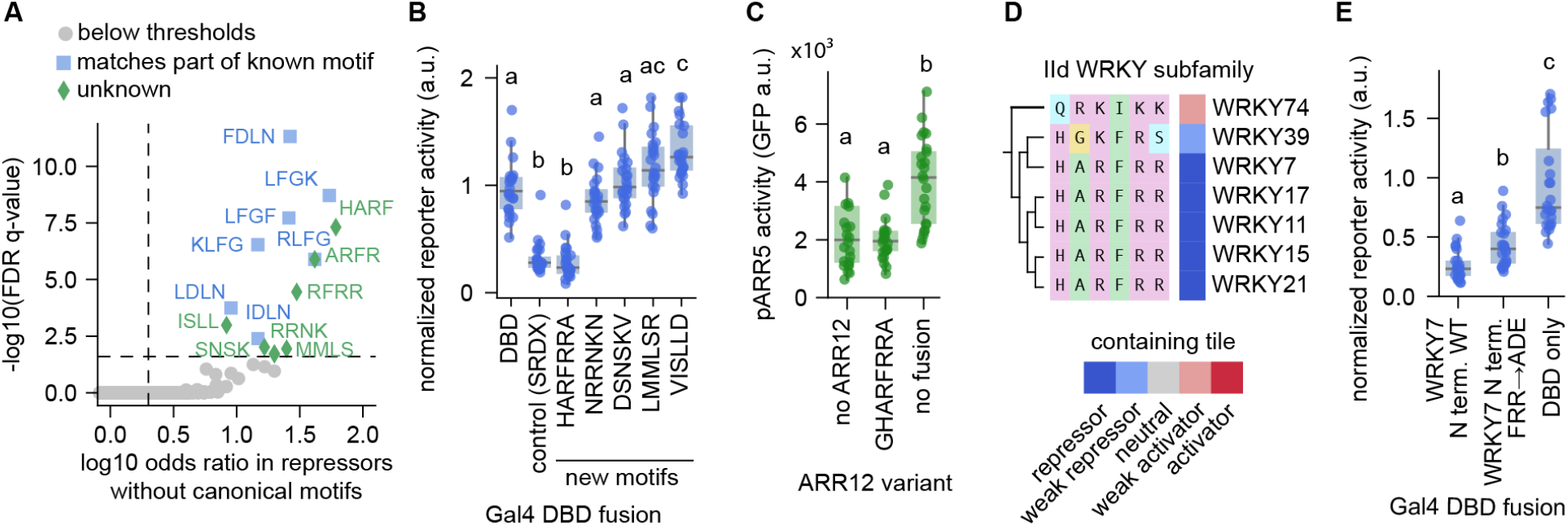
De novo motif discovery identifies a conserved WRKY-associated repression peptide. **(A)** Volcano plot of 4-mers enriched in repressor tiles without canonical motifs. **(B)** Gal4 DBD fluorescence reporter assay in *N. benthamiana* leaves. Peptides were fused including ±1 flanking residue from their strongest parent RD. **(C)** Fluorescence reporter assay in *N. benthamiana* leaves where the Arabidopsis ARR12 TF activates expression of the Arabidopsis ARR5 promoter driving GFP. Fusing a candidate motif peptide to ARR12 abrogates activation. **(D)** Sequence-similarity dendrogram of the WRKY TF subfamily IId showing the conserved HARFRR peptide. **(E)** Fluorescence reporter assay in *N. benthamiana* leaves where the N-terminal regulatory region of WRKY7 (amino acids 1-257) is recruited to a reporter via the Gal4 DBD. Mutation of the FRR residues in the HARFRR region of the WRKY7 regulatory domain weakens its repression strength. Box plots in (B), (C), and (E) show the median ± IQR, whiskers correspond to minimum and maximum values excluding outliers. Letters on top of boxplots in (B), (C), and (E) indicate statistically different groups (P < 0.05 based on a Tukey post hoc honestly significant difference test), N = 24 (4 plants with 6 leaf discs per plant).

### Charged, aliphatic-rich sequence features characterize motifless repression domains

Motifs are pliable sequence elements whose quantitative activity is strongly modulated by their context, and at the same time, most RDs lack detectable conserved motifs altogether. To unify these observations, we hypothesized that canonical motifs represent particularly conserved and easy to identify realizations of a continuous repression sequence-to-function landscape shaped by flexible and distributed sequence features (*58*). The hypothesis makes several predictions. First, motifless RDs should be predictable based on their biochemical and biophysical properties. Second, motif-containing and motifless RDs should share these properties. Finally, this implies that this putative repression-associated chemistry should remain detectable in motif-containing RDs even after their canonical motifs are artificially removed.

To test these predictions, we trained a random forest classifier based on the same set of biochemical and biophysical sequence features we used for ADs and tasked it with separating RDs from non-RDs. To remove the effect of motifs, we first performed in silico mutagenesis across the whole library, swapping consensus residues of known motifs with random residues, independent of tile activity (Fig. 5A). SHAP analysis indicated that the most important features for classifying tiles as RDs were an enrichment of basic residues, high density of charged residues, an enrichment of hydrophobic residues (in particular isoleucine), and an abundance of di- and tri-peptides containing positive and hydrophobic amino acids (Fig. 5B). Removing motif-containing tiles altogether produced similar results with similarly good predictive performance and recovered the same major predictive features (Fig. S17A,B) (Table S3). Hence, motifless RDs are predictable from biochemical and sequence features. Even after in silico disruption of their canonical motifs, motif-containing RDs retained substantially higher prediction scores than mutated motif-containing non-repressors, although their scores were lower than those of naturally motifless RDs (Fig. 5C). Thus, repressor tiles are biased toward a repression-associated chemistry which is shared between motif-containing and motifless RDs. Interestingly, many of the di- and tri-peptides enriched in repressors were also enriched among yeast-specific ADs in our cross-host analysis, even though by and large these sequences are not RDs (Fig. 2H, Fig. S17C-F).

**Figure 5.**
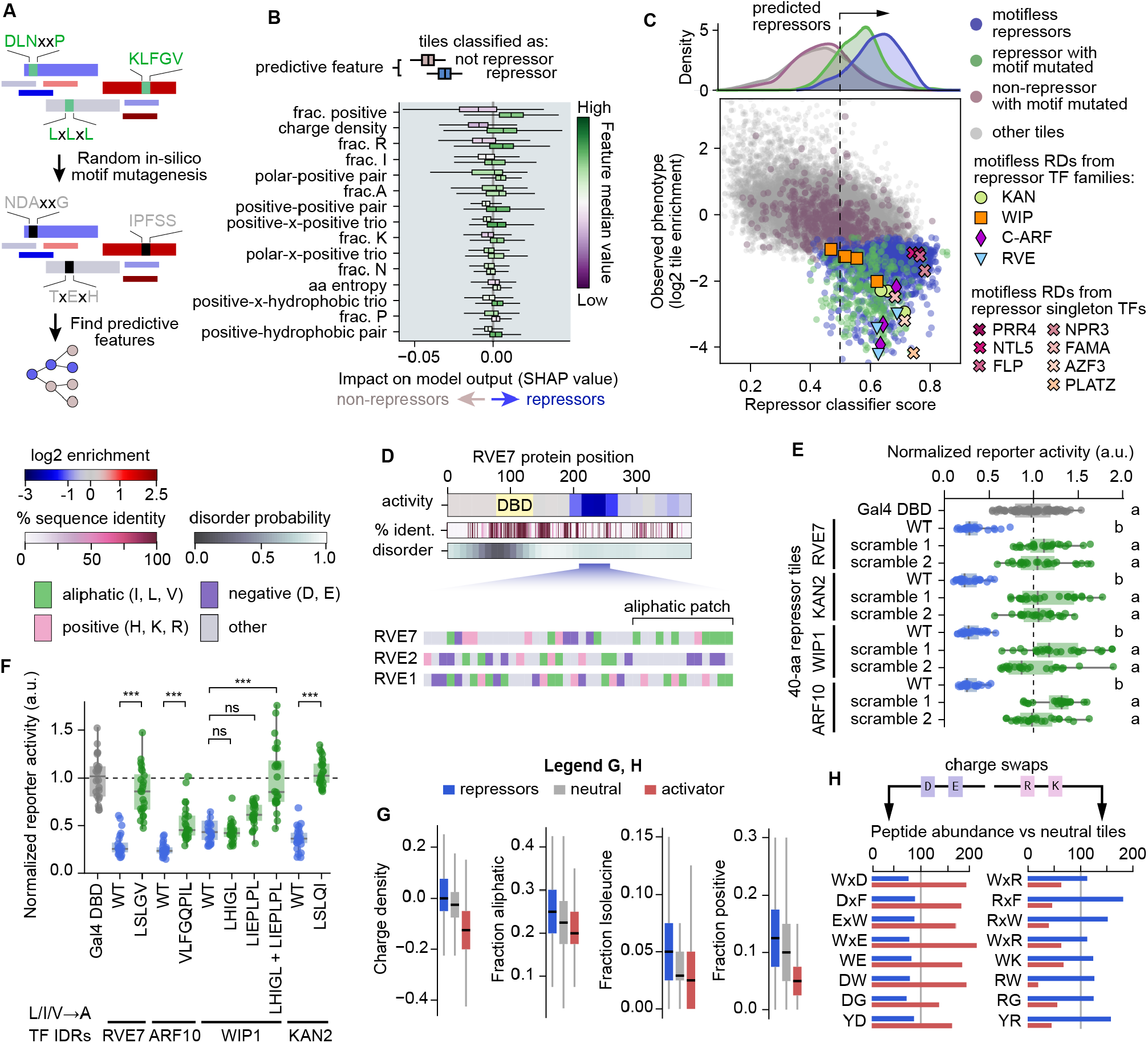
A shared distributed grammar unifies repression domains. **(A)** Schematic of the strategy to identify repression-associated sequence features independently of motifs. Canonical-motif consensus residues were randomly mutated in silico before training a random forest to distinguish RDs from non-RDs, forcing prediction to rely on motif-independent sequence features. **(B)** Summary of the most predictive sequence features. Shown is the mean ± SD SHAP value for the top 15 most discriminating features between non-repressors (left) and repressors (right). Features that are predictive when high are shown in yellow, in purple are features that are predictive when low. **(C)** Top: density distribution of classifier values for 4 classes of tiles in the library. Even though their motifs were in silico mutated, motif-containing RDs have higher scores than motif-containing non-RDs. Bottom: Scatterplot of tile activity (ENTRAP-seq enrichment, y-axis) versus classifier score (x-axis). Highlighted are motifless RDs from known repressor TF homologs, or singleton TFs (without a motifless repressor homolog). **(D)** Example of a protein family containing RDs with high prediction scores, RVE1, 2, and 7. These RDs are usually located in densely charged tiles with an overall positive charge and contain clusters of L, I, or V residues (aliphatic patches). **(E)** Fluorescence reporter assay testing 40-aa motifless RD tiles from the proteins highlighted in (C) and their scrambled versions preserving amino acid composition. Letters on top of boxplots indicate statistically different groups (P < 0.05 based on a Tukey post hoc honestly significant difference test). **(F)** Fluorescence reporter assay of the activity of the full IDR of motifless repressors highlighted in (C). Repression strength was tested upon mutating patches of L, I, or V residues to A. *** p<0.001, ** p< 0.01, * p<0.05, ns not significant using a two-sided Welch’s t-test. **(G)** Boxplots showing features predictive of repression that vary in the opposite direction in activators with respect to neutral tiles. **(H)** Percent abundance of charged di- and tri-peptides with respect to neutral tiles. Peptides on the right correspond to charge-swapped versions of corresponding peptides on the left. Box plots in (E)-(G) show the median ± IQR, whiskers correspond to minimum and maximum values excluding outliers. In (E) and (F) N = 24 (4 plants with 6 leaf discs per plant).

To identify higher-order sequence patterns associated with motifless repression domains, we combined model predictions with evolutionary conservation and functional evidence from the literature. We focused on strongly repressive, high-scoring tiles lacking known repression motifs and selected those derived from TFs previously characterized as repressors (Fig. 5C). We reasoned that sequence features contributing to motifless repression should be apparent among paralogous RDs that retain repressor activity. We therefore used sequence alignments to identify corresponding regions among paralogous TFs and kept those that were also high-scoring motifless RDs. This approach identified sets of paralogous motifless RDs across several subfamilies with evidence for repressive activity, including KANADI (KAN1–4) (*63*), REVEILLE (RVE1/2/7) (*64*, *65*), class C ARFs (ARF10/16/17) (*66*, *67*), and WIP1/TRANSPARENT TESTA1, 3, 4, and 5 (*68*) (Fig. 5C). Visual inspection of these motifless RDs revealed several architectural features that recur across subfamilies and among paralogs within each subfamily (Fig. 5D, Fig. S18A-C). These domains generally exhibit weak primary-sequence conservation and are frequently located within predicted IDRs. Compositionally, they often contain aliphatic patches—short segments of ∼5–10 residues enriched in L, I, and V—embedded within regions with a high density of charged residues, particularly positive ones (Fig. 5D, Fig. S18A-C). We identified similar features in RDs of previously characterized repressors without corresponding RDs in their paralogs (e.g., FAMA), or whose paralogous RDs instead contained canonical motifs (e.g., PRR4 and AZF3) (Fig. S18D-F).

From a high-level conceptual perspective, these motifless RDs are reminiscent of ADs in their weak primary-sequence constraints, frequent localization to IDRs, and apparent reliance on distributed features. If, as in many acidic ADs, activity is encoded mostly by amino acid composition rather than residue order, these RDs should retain activity upon scrambling their residues (*19*, *20*, *41*). We compared 3 motifless RDs with two of their sequence-scrambled counterparts, none of which acted as repressors (Fig. 5E). Thus, although these RDs lack canonical motifs and are poorly conserved, chemical composition alone is not sufficient to encode repression and the non-random ordering of residues must carry critical information for the activity of these sequences (Fig. S18G). Aliphatic L, I, and V residues are often part of canonical and degenerate functional repression motifs (Fig. 3), and have high predictive weights in our repressor-specific model, as well as our general plant activity regressor (Fig. 5B, Fig. S13, Fig. S17). These residues are also enriched in yeast-specific activators compared to plant activators (Fig. 2H). Therefore, we hypothesized that patches of these aliphatic residues might contribute to a plant repression grammar. To test the functional contribution of these aliphatic patches, we mutated clusters of L, I, and V residues to A within their native IDR context.

Gal4-DBD fusions to these IDRs acted as repressors, and aliphatic patch mutations weakened their activity to varying degrees (Fig. 5F). The same was true for the full-length RVE7 protein repressing one of its target promoters in a transient assay (Fig. S18H-J) (*64*). These results support a model where motifless RDs depend in part on the local organization of aliphatic residues embedded in a supportive biochemical context, a context that is shared with RDs carrying conserved canonical motifs (*58*). Taken together, our dissection suggests that plant repression can retain a greater dependence on residue patterning than the largely composition-driven activation grammars described for many ADs. This greater dependence on residue ordering or grammar may explain why short linear motifs have remained such a prominent feature of repression, but not activation, in plant TFs.

Compared with neutral tiles, many of the features associated with repression varied in the opposite direction from those associated with activation (Fig. 5G,H). This contrast was particularly apparent for charge-associated features: RDs were more positively charged than neutral tiles, which were in turn more positively charged than ADs (Fig. 5G). This polarity extended beyond charged residues to other amino acids, particularly aliphatics such as I and L, and was also evident in local sequence organization (Fig. 5H). Di- and tri-peptides juxtaposing positive and hydrophobic residues were enriched among RDs and depleted among ADs, whereas analogous peptides swapping negative for positive residues showed the opposite association. Hence, ADs and RDs are organized in opposite directions along several biochemical dimensions, most saliently those involving charge.

## Discussion

Sequencing technologies now routinely sample the enormous sequence diversity of life, but understanding the sequence-to-function mapping of this genomic diversity has lagged behind. Functional biology at the genomic scale still depends on a small number of tractable model systems such as yeast, where experimenting on thousands of sequences is feasible. This experimental tractability often comes with a privileged epistemological status: mechanisms first dissected in yeast can come to stand for general eukaryotic principles when there is no a priori reason to assume that they should. Transcriptional effector domains exemplify this bias: large-scale studies have defined activation mostly in yeast and animal cell culture, and repression largely in human cell culture. Our study demonstrates how functional genomics tools in plants, such as ENTRAP-seq, can extend genome-scale sequence-to-function measurements beyond opisthokont model systems. This is more than a philosophically principled position: our findings show how using host diversity as a comparative experimental lever can reveal evolutionary principles that remain idiosyncratic findings when studied in a single organism.

Plants contain numerous TF families that arose and diversified within Archaeplastida, many of them before the emergence of land plants (Fig. S19) (*69*). The abundance of shared ADs within many of these green lineage and land plant TF families is hard to explain only by these regulatory domains having been inherited as sequence modules from older eukaryotic ancestors. Instead, given the simplicity of many of these sequence features, our results suggest that the conserved acidic activation grammar can repeatedly and convergently emerge during plant evolution and need not be inherited as a conserved sequence module from more ancient protein lineages (*70*).

Conceptually as well as chemically, plant activation and repression domains appear to be governed by comparable principles. Both are frequently localized to IDRs, exhibit low primary-sequence conservation, and can be predicted from amino acid composition and biophysical features. Remarkably, many of their defining chemical features display a polarity around the acidic-to-basic axis, which tracks their contrasting activity. This chemical polarity raises the possibility that the biochemical basis of activation and repression in plants is itself intrinsically antagonistic. In this picture, the transcriptional effect of a protein sequence emerges from competing activation- and repression-biasing features, whose balance defines a continuum of regulatory potentials. This view may also help explain yeast-specific ADs: these sequences carry weaker activation-promoting features than shared ADs, while simultaneously being enriched in aliphatic residues such as I, L and V that collectively bias sequences toward repression in plant cells.

By bringing the sequence-function landscape of an entire plant TF repertoire within experimental reach, this work establishes a foundation for moving from description of regulatory domains to prediction and design. However, prediction from sequence is only a first step. We still do not fully understand how canonical acidic ADs work mechanistically, let alone the broader regulatory classes of ADs and RDs uncovered here, making it essential to connect sequence grammars to the molecular pathways and interaction partners that give rise to them.

## Acknowledgements

We are very thankful to a number of colleagues who were open to discussing our results. Their insights greatly contributed to better representing the data as well as to contextualize it within the broader literature. Alexander Leydon helped us better understand the function of the plant repression machinery in yeast cells and the functional role of sequence variation in plant repression motifs. Max Staller sharpened our thinking about differences between yeast and animal activation domains and gave us critical feedback for data visualization. Claire LeBlanc helped us to think more critically about predictive models of activation domain function. Lucas Waldburger provided feedback on the design architecture of machine learning models of activation domains and shared his ENTRAP-seq cartoon for this publication. We are also thankful to the members of the Shih lab, especially Liam Kirkpatrick, who graciously agreed to all of our requests for yeast and plasmid DNA. Next generation sequencing was supported by QB3 Genomics, UC Berkeley, Berkeley, CA, RRID:SCR_022170. This material is based upon work at the Joint BioEnergy Institute (JBEI) supported by the U.S. Department of Energy, Office of Science, Biological and Environmental Research Program under contract DE-AC02-05CH11231 with Lawrence Berkeley National Laboratory. A.E.S and M.A.K were also supported by the National Institute of General Medical Sciences of the National Institutes of Health under Award Number 5T32GM132022. I.O. is funded by a University of California, Berkeley Mentored Research Award, Graduate Research Fellowship from the National Science Foundation, and a Gilliam Fellowship from the Howard Hughes Medical Institute. J.K.N. acknowledges funding for this work from Biohub San Francisco, Bakar Fellows Program, Pew Scholars Program. This research used the Lawrencium computational cluster resource provided by the IT Division at the Lawrence Berkeley National Laboratory (Supported by the Director, Office of Science, Office of Basic Energy Sciences, of the U.S. Department of Energy under Contract No. DE-AC02-05CH11231)

## Author contributions

P.M.S. provided funding, strategic project guidance, and advice on experiments and data analysis. S.A. and A.E.S performed all experiments, unless indicated. S.A., A.E.S., J.T.R., An.L., and J.T.B. performed data analysis. A.E.S., J.T.R., An.L., and J.T.B. designed and implemented predictive models. P.M.S., S.A., A.E.S., An.L., J.T.R., J.T.B., I.O., M.A.K., and Am.L. participated in project discussions and/or inception. I.O. performed experiments in human cell lines. H.G.M. provided materials, support and guidance to J.T.R., and J.T.B. J.K.N. provided materials, support and guidance to I.O. S.A. and P.M.S wrote the manuscript.

## Data and Materials Availability Statement

Code used for feature annotation and predictive models, as well as supplementary tables can be found in https://github.com/shih-lab/Arabidopsis-TF-library-public. Correspondence and requests for materials should be addressed to Patrick M. Shih.

## Disclosure of Generative AI Use

During preparation of this manuscript, the authors used OpenAI ChatGPT (GPT-5.6; accessed 3/15/2026-9/29/2026) throughout the manuscript to edit for grammar, clarity and conciseness. No factual claims or interpretations were generated by AI. No citations were chosen by AI. Generative AI did not determine the experimental design, scientific judgments, final interpretations or conclusions. The authors used Cursor (version 3.12.17, with Grok 5.6; accessed 3/15/2026-9/29/2026) to assist with implementing and adapting code from previously published software repositories and with organizing, cleaning, and debugging original code written by the authors for this study. All AI-assisted text and code were reviewed, revised, and tested by the authors. No protected health information or trade secrets were entered into these tools, and no unpublished research material was provided to non-approved external AI models. The authors take full responsibility for the accuracy, integrity, and originality of the final work.

## Materials and Methods

### Gene library design

The primary gene models of Arabidopsis genes annotated as transcription factors (TF, 1,918 genes) were obtained from the Araport11 genome assembly (version June 2020). Protein coding sequences were tiled starting from the N terminus using 40-aa windows with a 20 amino acid stride, perfectly matching a subset of the tile sequences tested in yeast by Morffy et al. The DBDs of 67.3% of proteins were obtained from the PROSITE database. The domains of the remaining proteins lacking an annotated DBD from PROSITE were manually checked for DNA binding association in the UNIPROT database. After manual curation, 14% of proteins did not have an annotated DBD. Any tiles overlapping annotated DBDs were excluded. As a result of this DBD-exclusion filter, 59 TFs consisting of little more than a DBD had no tiles left and were therefore not part of the library. C-terminal tiles of less than 40-aa were also excluded for a final total of 26,023 tiles. DNA chisel was used to codon-optimize sequences for *N. benthamiana* avoiding repetitive sequences and cut sites for common restriction enzymes, keeping the GC content between 45% and 65% while avoiding hairpins and homopolymeric repeats of 6 or more nucleotides (*71*). The resulting tiles were split into 12 sub-libraries following the order of their genomic coordinates (AGI locus number): 10 libraries of 1,900 tiles (ATF1-5 and ATF9-13), one sub-library of 3,123 tiles (ATF14), and one sub-library of 3,900 tiles (ATF6). Each sub-library also contained 100 shared internal control 40-aa tiles measured in previous studies and spanning a wide dynamic range of activity. Each sub-library was flanked by BsaI cut sites for cloning into the ENTRAP-seq backbone and a unique pair of primers to isolate it from the rest of the libraries by PCR. Sub-libraries were synthesized in bulk as oligo pools (Twist Bioscience).

### Library cloning into ENTRAP-seq vector

All ENTRAP-seq runs as well as fluorescence assays using Gal4 DBD fusions used plasmid backbone pSA676v6. The oligo pool containing all libraries was resuspended to 0.75 ng/µL of DNA in water. Each sublibrary was PCR amplified with unique primer pairs using 6 50-µL Q5 2X Mastermix reactions (NEB #M0492S) with 1 µL of oligo library as template and 15 cycles with a tm of 61°C. PCR products were purified using the zymo PCR cleanup kit (#D4029), washing twice with 600 µL of wash buffer and eluting in 30 µL of water. Amplified oligos were digested overnight with BsaI and purified again using the zymo PCR cleanup kit. The pSA676v6 backbone was digested with BsaI and rSAP overnight and run on an agarose gel to isolate the backbone from a BsaI-flanked GFP dropout cassette. Ligations were performed using 6.6 µL of T7 ligase (NEB #M0318S), 66 ng of backbone DNA, and 33 ng of library insert in 132 µL final volume, resulting in a 3:1 ratio of insert to backbone. Ligations were incubated using 3 cycles of 22°C for 2 hr and 16°C for 4 hr and purified using the zymo PCR cleanup kit by eluting in 15 µL of water.

### Library transformation into *E. coli* and *Agrobacterium*

E. coli electroporation reactions were prepared using 2 µL of purified ligation and 1 tube of 10-beta electrocompetent cells (NEB #C3020K). Cells were recovered for 1 hr in 1 ml of LB media shaking at 37C. After recovery, 1 µL of cells was diluted in 100 µL of LB for CFU counting in diagnostic plates. The rest of the culture was transferred to a flask containing 100 mL of LB with 50 µg/mL kanamycin and incubated overnight at 37°C. Libraries were used if the extrapolated number of CFUs after the 1-hr recovery was at least 10X the size of the library. Cryostocks and a plasmid miniprep were prepared from each overnight E. coli library culture. The library was transformed into *Agrobacterium* GV3101 scaling up the electroporation to obtain at least 10X times as many CFUs as the library size. For a typical library we used 5 electroporation mixes with 400 ng of library DNA, 5 µL of electrocompetent bacteria and 95 µL of water. Each electroporation was recovered in 1 ml of LB shaking for 2 hr at 30°C. Recovered cultures were pooled together and 1 µL was set aside to perform serial dilutions for CFU counting. Libraries were used if Agrobacterium CFU counts in the recovery volume were at least 10X the number of tiles in the library. Pooled cultures were transferred to a flask containing 100 ml of LB with 50 µg/mL kanamycin, 30 µg/mL gentamicin, and 100 µg/mL rifampicin. Liquid cultures were grown for ∼48hr shaking at 30°C, after which multiple cryostocks were prepared.

### Plant material and growth conditions

Wild-type *N. benthamiana* plants from the JBEI seed bank were grown for 5 weeks in Sungro Sunshine soil mix #4 (aggregate plus) in long-day conditions (16L:8D) at 25 °C in 60% humidity under 150 μmol m−2 s−1 photosynthetically active radiation. Plants were returned to the same conditions after agroinfiltration.

### Agroinfiltration

ENTRAP-seq library infiltrations and agroinfiltration for fluorescence reporter assays were prepared using the same protocol, unless otherwise stated below. Cryostocks were inoculated into LB containing 50 µg/mL kanamycin, 30 µg/mL gentamicin, and 100 µg/mL rifampicin and grown overnight shaking at 30°C. For gene libraries, 1-mL aliquot of a cryostock was inoculated in 10 mL of LB to ensure good representation. The day of infiltration, cultures were diluted 1:10 in the same growth media and incubated for 2-3 more hours in the same conditions until reaching an optical density at 600 nm (OD_600_) of approximately 1.0. Cultures were spun down at 4,000*g* for 10 minutes and resuspended in infiltration buffer (10 mM MgCl2, 10 mM MES, and 200 μM acetosyringone, pH 5.6) to a similar final volume. Cultures were incubated with gentle shaking at room temperature for 1 hr after which OD_600_ was measured to determine dilution factors. Gene libraries were infiltrated at OD_600_ 0.0025. For fluorescence reporter assays, each culture was diluted to OD_600_ 0.4 and cultures were mixed in equal ratios. Two of the 6th-8th true leaves to emerge were syringe infiltrated and marker was used to label infiltrated areas.

### Optimized ENTRAP-seq

Nuclei were isolated from infiltrated *N. benthamiana* tissue 3 days post infiltration using a protocol adapted from published INTACT procedures (*72*, *73*). Magnetic sorting of nuclei was performed using an optimized version of the original ENTRAP-seq protocol. All steps following tissue grinding were performed in a cold room or on ice unless otherwise indicated. Nuclei purification buffer (NPB) contained 20 mM MOPS (pH 7.0), 40 mM NaCl, 90 mM KCl, 2 mM EDTA, and 0.5 mM EGTA and was freshly supplemented with 0.5 mM spermidine, 0.2 mM spermine, protease inhibitor, and 1 mM PMSF immediately before sample preparation. NPBt consisted of NPB supplemented with 0.1% (v/v) Triton X-100. For each sample, 10 g of infiltrated tissue was cut with a razor blade into approximately 1x1 cm pieces. The tissue was submerged in 100 mL NPB and vacuum infiltrated for 5–10 min. The vacuum was released and infiltration was repeated until the tissue was thoroughly infiltrated. The tissue was then rinsed three times with tap water, blotted dry with paper towels, and flash-frozen in liquid nitrogen. Samples were either stored at −80°C or processed immediately. The tissue was ground to a fine powder in liquid nitrogen and resuspended in 100 mL ice-cold NPB. The suspension was passed through a 70-µm cell strainer and centrifuged at 500*g* for 10 min at 4°C. Nuclei pellets were gently resuspended and combined in 10 mL NPB, then passed through a 100-µm cell strainer to remove residual tissue debris. For magnetic purification, 30 µL Dynabeads (Thermo Scientific #65001) per sample were washed in NPB and resuspended in 10 mL NPB. The 10-mL bead suspension was combined with the 10-mL nuclei suspension and mixed for 30 min at 4°C with gentle continuous rotation. The resulting 20-mL suspension was divided between two 50-mL tubes and diluted to a total volume of 100 mL with NPBt. Bead-associated nuclei were collected on a magnetic rack for 30 min. The supernatant was transferred to fresh tubes, while the bead-associated nuclei were resuspended and combined into 50 mL NPBt and set aside. The supernatant was subjected to a second 20-min magnetic separation to recover residual bead-associated nuclei. A representative 50-mL portion of the new supernatant, comprising equal volumes from the two tubes, was retained as the unbound fraction. Bead-associated material recovered from the first and second magnetic separations was combined and subjected to three additional washes in NPBt, with 20-min magnetic collection after each wash. Nuclei were first collected magnetically for 20 min, followed by a second 50-mL NPBt wash and a final 15-mL NPBt wash. The purified bead-associated nuclei were finally resuspended in 3 mL NPB and were pelleted at 500 *g* for 10 min and resuspended in 250 µL CTAB extraction buffer. The retained unbound fraction was centrifuged at 500*g* for 10 min, resuspended once with NPB, and centrifuged again at 500 × *g* for 5 min. The resulting pellet was resuspended in 1 mL of cetyltrimethyl-ammonium bromide (CTAB) extraction buffer (100 mM Tris-Cl pH 8, 20 mM EDTA pH 8, 1.4 M NaCl, 2% (w/v) CTAB, 1% PVP 40,000). Both bead-associated and unbound fractions were stored at −20°C until DNA extraction.

### DNA extraction

DNA was purified using a CTAB–phenol extraction protocol. Samples stored in CTAB were incubated at 65°C for 30 min with vortexing every 5–10 min. An equal volume of phenol:chloroform:isoamyl alcohol (25:24:1, v/v/v) was added to each sample, and tubes were vortexed vigorously. Samples were centrifuged at 20,000*g* for 2 min at room temperature, and the upper aqueous phase was transferred to a new tube and mixed with an equal volume of CTAB. Samples were incubated at 65°C for 20 min with vortexing every 5–10 min. Chloroform:isoamyl alcohol (96:4, v/v) was then added at a volume equal to the volume of CTAB added in the previous step. Tubes were vortexed vigorously and centrifuged at 20,000 × g for 2 min at room temperature. The upper aqueous phase was transferred to a new tube and 0.7 volumes of isopropanol relative to the aqueous phase were added and mixed by gentle inversion. Samples were incubated at −20°C for 20 min, after which DNA was pelleted by centrifugation at 20,000 × g for 15 min at room temperature. Pellets were rinsed with 70% ethanol and air-dried on a bench top. Dry pellets were resuspended in 50 µL of water.

### DNA sequencing

Sequencing libraries were prepared using PCR with primers containing unique multiplexing indexes and Illumina adapter stubs (Table S5). A 150 µL Q5 (NEB #M0492S) PCR reaction was performed following the manufacturer’s guidelines using 3 µg of DNA as template, 61°C annealing, and 15 s of elongation for 25 cycles. Amplicons were purified by double-sided SPRI bead size selection and the size of the bead-bound DNA was verified using a fragment analyzer. Sequencing libraries were quantified by qPCR and pooled equimolarly. Libraries were sequenced using 100 bp single-end reads in the Illumina NextSeq 2000 platform with 30% PhiX. Sequencing depth was chosen so as to obtain approximately 10-20M raw unfiltered reads per 2,000-tile library. This depth ensures that only 5-10 tiles per library have fewer than 50 reads.

### Calculation of tile enrichment from sequencing

Reads were mapped back to library tiles using 2FAST2Q, using the 76 bp of tile-derived sequence recovered from the 100-bp single-end reads. A per-nucleotide Phred quality threshold of 30 was applied across the 76-bp sequence prior to mapping, and no mismatches were allowed. For each sequencing sample, the abundance of each tile was normalized by dividing the number of reads assigned to that tile by the total number of reads successfully mapped to any library tile in the same sample. Hence, normalized tile abundance represents the fraction of all mapped reads in a sample contributed by that tile. At this point, tiles with zero reads in either the pulldown or flowthrough sample were discarded, but no minimum read coverage filter was performed. Tile enrichment was calculated as the ratio of normalized tile abundance in the pulldown fraction to that in the flowthrough fraction.

### Harmonization of enrichment values across libraries

Tile enrichment was first log2 transformed. To harmonize enrichment measurements across independently assayed libraries, log2 enrichment values were robustly normalized using the 100 shared internal control tiles. For each library replicate, the distribution was centered on the median log2 enrichment of the validated transcriptionally neutral internal controls. Extreme outliers in the 100 control set were excluded using a z-score threshold of 4.5 among the control set. The spread of the remaining internal controls around the neutral-control median was estimated using the median absolute deviation (MAD), scaled by 1.4826 to obtain a robust estimate equivalent to SD. Normalized tile enrichment was calculated by subtracting the median log2 enrichment of the neutral internal controls from each tile’s log2 enrichment and dividing by the scaled MAD of the internal control set around the neutral-control median. Biological replicates were normalized independently and normalized log2 enrichment values were subsequently averaged for each tile.

### Tile activity classification

Tile activity classes were assigned separately for each library after averaging normalized enrichment across biological replicates. Tiles within ±1 SD of the mean activity of validated neutral controls were classified as neutral. Activator and repressor thresholds were set at ±1.25 SD from the mean of all tiles in that library after control-based normalization, using the more conservative boundary if this overlapped the neutral-control range. Tiles between the neutral and activator/repressor thresholds were classified as weak activators or weak repressors, respectively.

### Fluorescence reporter assays in N. benthamiana

Gal4 DBD fusions were expressed from the same vector backbone used for ENTRAP-seq (pSA676v6). In addition to the Gal4 DBD fusion and BirA, this vector contains an mCherry reporter driven by six tandem Gal4 UAS elements followed by a minimal 35S promoter. The mCherry reporter is linked by a T2A peptide to the GFP-tagged outer nuclear envelope protein used for ENTRAP-seq. Agrobacterium strains carrying the Gal4 DBD fusions were co-infiltrated with a normalization strain carrying a constitutively expressed BFP transgene driven by the pMAS promoter (pSA639). A construct in which a stop codon immediately followed the Gal4 DBD, rather than an effector-domain fusion, was used as the Gal4 DBD-only control. To increase the dynamic range of repressor assays, a third Agrobacterium strain was included carrying an mCherry reporter driven by a stronger promoter containing six tandem Gal4 UAS elements embedded within a leaky synthetic promoter (pSA521v3). Thus, mCherry fluorescence in repressor assays reflected the combined signal from the reporter encoded by pSA676v6 and the additional reporter encoded by pSA521v3. Each strain combination was infiltrated into two leaves per plant across four plants, with two to four strain combinations infiltrated at spatially separated positions on each leaf. Three days after infiltration, three 6-mm leaf discs were collected from each infiltrated region and placed abaxial side up in 350 µL tap water in a 96-well plate. mCherry fluorescence (587-nm excitation, 615-nm emission) and BFP fluorescence (400-nm excitation, 454-nm emission) were measured using a Synergy 4 microplate reader (BioTek), with gain set to 100 and read height to 10.5 mm. Reporter fluorescence for each leaf disc was first adjusted to BFP by calculating the mCherry/BFP fluorescence ratio. Normalized reporter activity was then calculated by dividing this ratio by the mean mCherry/BFP ratio of leaf discs infiltrated with the Gal4 DBD-only control.

### Plant reporter-assay statistics

Each construct was tested in eight independently infiltrated leaf regions distributed across two leaves on each of four plants. Three leaf discs were collected from each infiltrated region, yielding 24 measurements per construct. Each disc was treated as an individual sample for statistical analysis.

### Plant–yeast comparison

Arabidopsis tiles captured by ENTRAP-seq were matched to the corresponding tiles measured in yeast by Morffy et al. Only tiles with measurements in both datasets were included in cross-host comparisons. Tiles were classified as activators in yeast if they had a PADI score ≥ 1, as defined by the authors. Tiles were classified as activators in plants according to the criteria listed in the main text, namely, if they had a log2 enrichment greater than 1.25 SD from the mean of the distribution of two averaged biological replicates. Tiles that satisfied both criteria were classified as shared activators. Yeast-specific activators correspond to tiles with PADI scores ≥ 1 and belonging to the ENTRAP-seq categories neutral, weak repressor, and repressor. Tiles classified as neutral, weak repressor, and repressor by ENTRAP-seq with PADI score < 1 were considered not activators. Note that tiles falling into the weak activator ENTRAP-seq category are not assigned to any category.

### Fluorescence reporter assays in human cells

Activities of dCas9-fusions were measured by transient transfection and recruitment to the endogenous EPCAM gene. HEK293T cells were maintained under standard culture conditions (DMEM supplemented with 10% (v/v) FBS (VWR), 100 U/mL penicillin and 100 µg/mL streptomycin) and seeded into 96-well plates at 15,000 cells per well one day prior to transfection to enable biological duplicate measurements for each dCas9-fusion. Cells were co-transfected with 150 ng of dCas9-effector plasmid and 75 ng of 4-pooled sgRNA plasmids targeting the endogenous EPCAM locus. The dCas9-effector plasmid consists of catalytically inactive Cas9 (dCas9), an HA tag, nuclear localization sequence, blue fluorescent protein (BFP), and the indicated effector domain. sgRNA expression plasmids contain an mCherry reporter to enable identification of sgRNA-positive cells. Two days post-transfection, cells were harvested and stained for surface expression of commercially available FITC labeled antibodies for EpCAM (BioLegend Cat. #324203) following the manufacturer’s protocol. Cells were analyzed by flow cytometry using BD FACSymphony A1 Cell Analyzer (BD Biosciences) flow cytometer. BFP, mCherry, and FITC-channel fluorescence were measured to identify cells expressing the dCas9-effector fusion, sgRNA, and target protein expression, respectively. All flow cytometry data were analyzed using FlowJo. Cells were first gated for single, live cells, followed by selection of BFP-positive and mCherry-positive populations corresponding to successful delivery of both dCas9-effector and sgRNA components. Effector activity was quantified by measuring target gene activation within the BFP-positive/mCherry-positive population using FITC-channel mean fluorescence intensity (MFI). The transcriptional activation domain VP16 was included as a positive control.

### Predictive features annotation

Tile sequence was used to calculate biophysical, compositional and patterning features, listed in Supplementary Table 1. Biophysical features included protein disorder calculated using Metapredict v3 (*74*), overall charge, charge segregation, hydrophobicity, and helical propensity. Compositional features included the frequency of each amino acid and the frequency of each amino acid class. Patterning features included the counts of dipeptides and gapped dipeptides of all possible combinations of amino acids and amino acid classes. Patterning features also included the degree of clustering of each amino acid and amino acid class, as well as the length of the longest homopolymeric run of each amino acid. The code used to annotate features can be found in https://github.com/shih-lab/Arabidopsis-TF-library-public/tree/main/other.

### Random forest classification of AD classes

Each tile was represented using a set of ∼900 sequence-derived features (see: Predictive features annotation). Random forest classifiers were then trained using five-fold stratified cross-validation. Each fold held out about one-fifth of the tiles and tiles from the same TF were assigned to either train or test to avoid data leak. Every tile received exactly one out-of-fold prediction from a model that had not seen it. Folds were stratified so that the class balance in the training and test loci stayed close to the full comparison. Within each fold we performed a feature selection step as follows: features were ranked by random forest impurity-based importance, and the 100 highest-ranking features were retained. Forests used 200 trees, a maximum depth of 10, a minimum of 15 samples to split a node, a minimum of 5 samples per leaf, and balanced class weights, so the larger class did not dominate. Separate classifiers were trained for plant-specific versus shared, yeast-specific versus plant-specific, and yeast-specific versus shared activators. Feature contributions to the five-fold classification forests were summarized with SHAP (SHapley Additive exPlanations)(*75*). For each fold, SHAP values for the positive class were computed on a random sample of 200 held-out tiles (1,000 tiles in total). Classifier performance is reported in Table S3.

### XGBoost regressor ensemble classification of AD classes

Each tile was represented using a set of ∼900 sequence-derived features (see: Predictive features annotation). Features and instances containing NaN feature values or missing observed values for plant enrichment or PADI score were removed to avoid misleading outcomes. Data were split into five cross-validation folds.. For each fold, an ensemble of XGBoost regressors (*76*) were trained on individual measurements (n=2) on the training data set and tested for predictive power on the mean values of the validation set. For each fold, Parameters for XGBoost regression, specifically the number of trees, depth of trees, and subsampling ratios for instances and features, were selected using scikit-learn’s (*77*) GridSearchCV with an inner 5-fold cross-validation on the current cross-validation fold’s training data. To limit the potential for data overfitting, five XGBoost regressors were trained using the selected parameters, matching the five regressors per cross-validation fold trained during the GridSearchCV. These five regressors were then pooled into an ensemble such that the ensemble prediction is the mean of the individual regressor predictions. Finally, the ensemble regressor was used as a binary classifier by assigning activator/non-activator labels according to predictions being greater/lesser than a threshold value. Feature contributions in the final ensemble regressor were evaluated using SHAP (SHapley Additive exPlanations) (*75*), pooling the calculated SHAP values for out-of-fold predictions on all 5 cross-validation folds to get one combined ranking.

### Random forest classification of motifless repressors

Random forest classifiers were implemented using scikit-learn (*77*), following the algorithm of Breiman (*78*). Because non-repressor tiles outnumbered repressors by ∼13:1, random forest models were trained using repeated class-balanced batches. In each batch, the complete set of eligible repressors was retained and an equal number of motifless non-repressor tiles was randomly sampled from the larger non-repressor population. The total number of batches was therefore the number of nonrepressors divided by the number of repressors. Features were first pre-selected by training a random forest on all repressors and an equal number of randomly sampled non-repressors and keeping the 100 features with the highest impurity-based importance. Each batch was split 80:20 training:testing with stratification to keep the 1:1 class balance. Tiles from the same TF were assigned to either train or test to avoid data leak. An independent random forest classifier was trained for each batch and each batch’s model was then used to score every tile in the dataset. A tile’s final score was the mean predicted probability of being a repressor across all batch models. Model performance was evaluated on a randomly held-out 20% of each batch and averaged across batches. Feature importance measured as SHAP values was calculated for this held-out 20% and averaged across batches. Classifier performance metrics are found in Table S3.

### Predictive models software: FINCHES

FINCHES: To predict binding propensity between activating tiles and Mediator surfaces, we used FINCHES (*40*) with the CALVADOS v2 interaction model (*79*). For each tile, we computed surface-patch interactions against a folded-domain target using `FoldedDomain.calculate_idr_surface_patch_interactions()` (12 Å default patch radius). Two folded Mediator subunit surfaces were used as binding targets: the KIX domain of *Arabidopsis thaliana* MED15a (residues 6–101, AlphaFold model) and activation-domain-binding domain 1 (ADBD1) of *Saccharomyces cerevisiae* Gal11/Med15 (residues 158–238, from the NMR structure PDB 2LPB, as used in (*40*)). The *E. coli* RNA polymerase alpha subunit C-terminal domain (alphaCTD; PDB 3K4G) was used as a negative control for binding. For each tile, the interaction matrix was reduced to attractive epsilon (`att_eps`), defined as the sum of negative (favorable) matrix elements. att_eps distributions for each AD group were compared to the neutral group within each target domain using Cliff’s delta.

### Predictive models software: ADPred

For prediction of acidic activation domains we used ADpred (implementation and pretrained models from the Fred Hutch repository, https://github.com/FredHutch/adpred, also available at https://adpred.fredhutch.org/) (*19*). ADpred integrates each input peptide’s amino-acid sequence with PSIPRED-predicted secondary structure and returns a per-residue probability (0–1) that each position lies in an activation domain. Our Arabidopsis TF library fragments are 40-aa tiles. ADpred is typically applied to 30 aa inputs, so for each 40-aa tile we extracted three overlapping 30 aa subsequences starting at 0, 5, and 10 aa within the tile (5 aa step). Each 30 mer was passed to ADpred.predict(); for that window we retained the maximum ADpred probability over scored positions. The ADpred score assigned to each tile was the maximum of those three window scores.

### Predictive models software: PADDLE

For prediction of acidic ADs using PADDLE we used pretrained model weights and inference code obtained from the authors’ public repository (https://github.com/asanborn/PADDLE) (*20*). The sequence-only PADDLE-noSS models were used. PADDLE predicts a Z-score for fixed 53-amino-acid input windows, but our Arabidopsis TF library used 40-aa tiles. To reconcile the length difference, we scored full-length Arabidopsis TF protein sequences by tiling them into 53-aa windows with a step size of 1 aa, and each window was scored. For each library 40-aa fragment, we then identified all scored 53-aa windows that contained that 40-aa tile and used the maximum PADDLE score as the PADDLE score assigned to that tile.

### Predictive models software: TADA

Activation-domain propensity was predicted for each tile using TADA (*4*), as implemented in the TensorFlow 2-compatible package TADA_T2 (https://github.com/ryanemenecker/TADA_T2). Each 40-aa sequence was scored directly, without padding or overlapping windows, giving one TADA score per tile.

### Predictive models software: ADHunter

Transactivation potential was predicted using ADhunter, a ResNet ensemble of 20 models trained to predict activation domain activity from per-residue protein-language-model embeddings (*24*). Input peptide sequences were embedded with ESM2 (`esm2_t33_650M_UR50D`; (*80*)), and each embedding was scored by the full 20-model ensemble. The reported prediction is the ensemble mean. Predictions were generated using the ADhunter command-line tool (`adhunter run --fasta-file <INPUT> --output-file <OUTPUT> --use-gpu`).

### Repression motif identification

Canonical repression motifs were identified by scanning each 40-aa tile for matches to the regular expressions listed in Supplementary Table 3. Tiles containing at least one match were classified as motif-containing, all others were classified as motifless.

### Regression models of repression-motif activity

For each motif-containing tile, we extracted a fixed sequence window spanning the motif core and its flanks and sought to predict tile log2 enrichment from the amino acid identities at nonconsensus (variable) positions in that window. Sequences were encoded with two classes of categorical features: linear (main-effect) features for a given amino acid at each variable position, and pairwise interaction features for amino acid pairs at every combination of two variable positions. Features were one-hot encoded and used to fit an L2-regularized (Ridge) linear regression with intercept. Raw Ridge predictions were then affine-calibrated on training data only by shifting along the y axis by a factor b and adjusting the slope by a factor of m. The same m and b were applied to held-out test tiles. In repression-weight logos, letter height at each position reports that amino acid’s calibrated main-effect (no pairwise interactions) contribution to predicted negative enrichment (mean-centered within position), i.e. its average additive effect at that site after accounting for the other fitted terms; pairwise coefficients are not shown in the logo.

### De novo repression 4-mer enrichment and candidate selection

First, tiles with any canonical repression motifs as listed in Supplementary Table 3 were removed from the dataset, regardless of their activity. Foreground was defined as tiles classified as repressors, the background corresponded to all other qualitative categories combined. All possible contiguous 4-mers were obtained from the background and foreground. Each 4-mer counted at most once per tile and only 4-mers with 10 or more occurrences were kept. Motif enrichment in the repressor foreground was calculated using a Fisher’s exact test. P-values were adjusted across all tested 4-mers using the Benjamini–Hochberg false discovery rate (FDR). Candidate motifs were required to meet FDR q ≤ 0.05 and odds ratio ≥ 2 (enrichment in repressors vs the rest).

### In silico motif mutagenesis

Canonical repression motifs as defined in Supplementary Table 3 are composed of consensus positions (e.g. D, L, N, and P in the DLNxxP motif) and free variable positions (e.g. the ‘x’ positions in DLNxxP). Each consensus position was mutated to a random aa drawn from a homogeneous distribution excluding the original aa. If the mutation resulted in the creation of a new canonical motif, a new random aa draw was performed. Variable ‘x’ positions were left untouched.

### Protein sequence alignments for sequence similarity trees

Sequences were aligned with MAFFT v7.526 using the --auto option, which chooses the alignment strategy based on dataset size (*81*). Before tree inference, poorly aligned and highly variable columns were masked. A column was kept only if at least 50% of sequences had a residue there, and the most common residue made up at least 50% of those non-gap residues. For each pair of sequences, we computed an uncorrected p-distance on the masked alignment: the fraction of mismatched positions, counting only positions where neither sequence has a gap. Trees were built from the resulting distance matrix by average-linkage hierarchical clustering (UPGMA) with SciPy v1.18.1 (scipy.cluster.hierarchy.linkage, method="average") in Python 3.13, and exported in Newick format (*82*). Tree leaf order was used to order rows in the family-level heatmaps and alignment figures.

## Supplementary Materials

### Supplementary Tables

Tables and scripts can be found at https://github.com/shih-lab/Arabidopsis-TF-library-public

Table S1. Tile sequences

Table S2. Protein features used for training models

Table S3. Performance metrics of random forest and regression-based classifiers

Table S4. List of canonical repression motifs and their defining regexes

Table S5. NGS primers

### Supplementary Figures

**Supplementary Figure 1.**
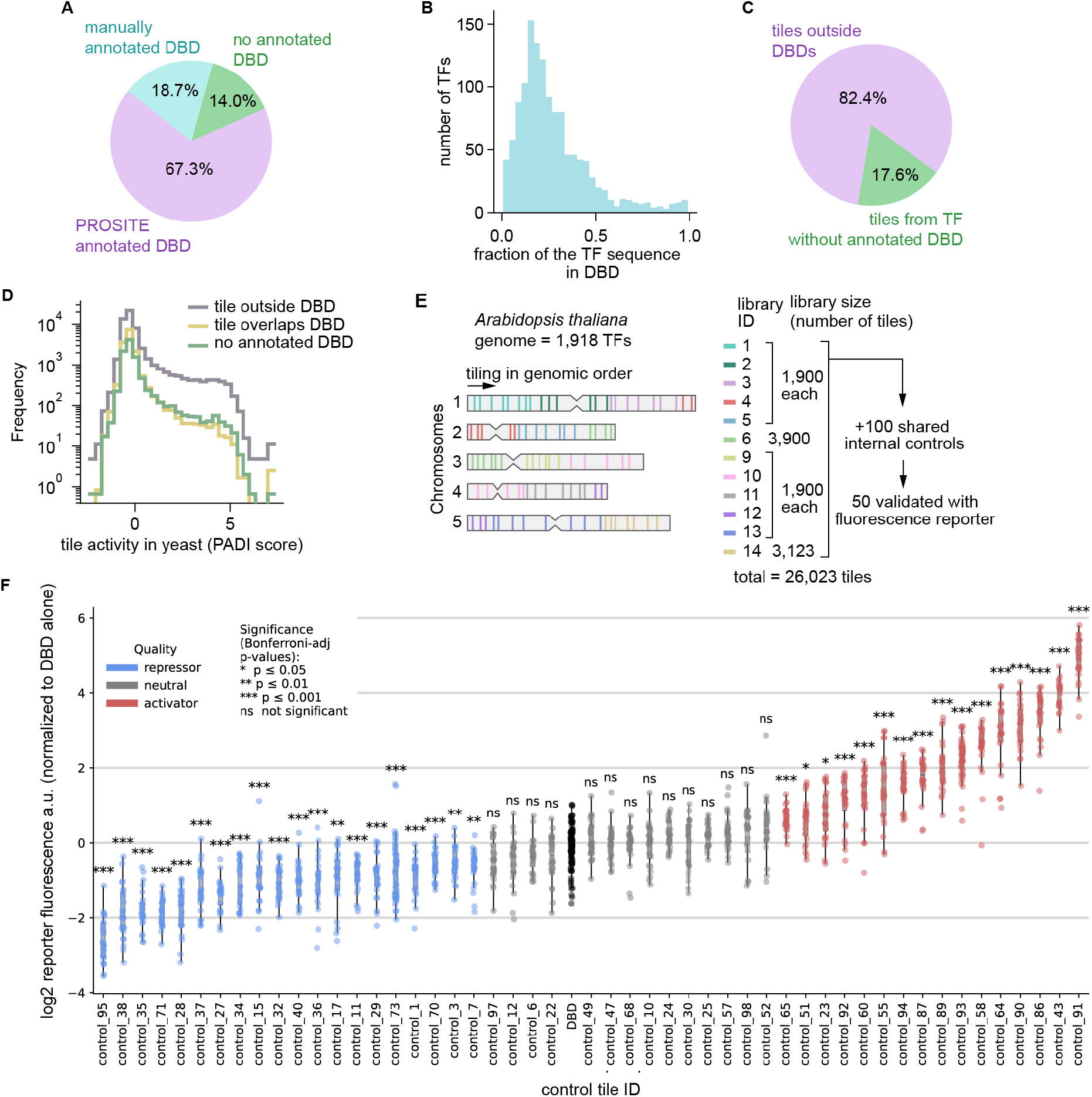
Details on library selection and internal controls. **(A)** Fraction of Arabidopsis thaliana TFs for which a DNA binding domain (DBD) annotation was identified. Out of 1918 TFs, 1290 (67.3%) had a domain annotated as DBD in the PROSITE database. The annotated protein domains in TFs without PROSITE DBDs (32.7%) were manually scanned for DBDs, resulting in the DBD annotation of 358 TFs (18.7%). The remaining 270 TFs (14%) were tiled in their entirety as no DBD annotation was obtained. **(B)** Histogram showing the fraction of the TF linear protein sequence corresponding to the DBD in TFs with annotated DBDs. **(C)** Fraction of library tiles belonging to TFs with or without DBD annotations. **(D)** Histograms showing the transcriptional activity of tiles from Arabidopsis TFs in *S. cerevisiae* (yeast). Tiles outside DBDs are enriched with activators compared to tiles overlapping DBDs. **(E)** Library construction approach. TF loci were selected in the order of their genomic coordinates. TF tiles were split into 12 sub-libraries. Each sub-library contained a common shared set of 100 internal controls, 50 of which were individually validated. **(F)** Validation of 50 shared internal controls using a fluorescence reporter assay. Internal control tiles spanning a wide dynamic range of activity were individually cloned as Gal4 DBD fusions in the ENTRAP-seq reporter construct (pSA676v6) and individually tested in *N. benthamiana* using a fluorescence reporter assay. The basal fluorescence of pSA676v6 is too low to resolve repression using fluorescence. Thus, tiles that were not significantly different from the Gal4-DBD control were subsequently tested for repression by coinfiltrating them with a reporter with a higher basal expression level consisting of 6xUAS sites upstream of the 35S promoter (pSA521v3). If tiles were significantly lower than Gal4-DBD in this second repression experiment, they were classified as repressors. A Bonferroni-corrected two-sided Welch’s t-test against the Gal4-DBD control was used to classify as repressor, neutral, or activators with *** p<0.001, ** p< 0.01, * p<0.05, ns not significantly different from Gal4-DBD. Box plots show the median ± IQR, whiskers correspond to minimum and maximum values excluding outliers. N = 24 (4 plants with 6 leaf discs per plant).

**Supplementary Figure 2.**
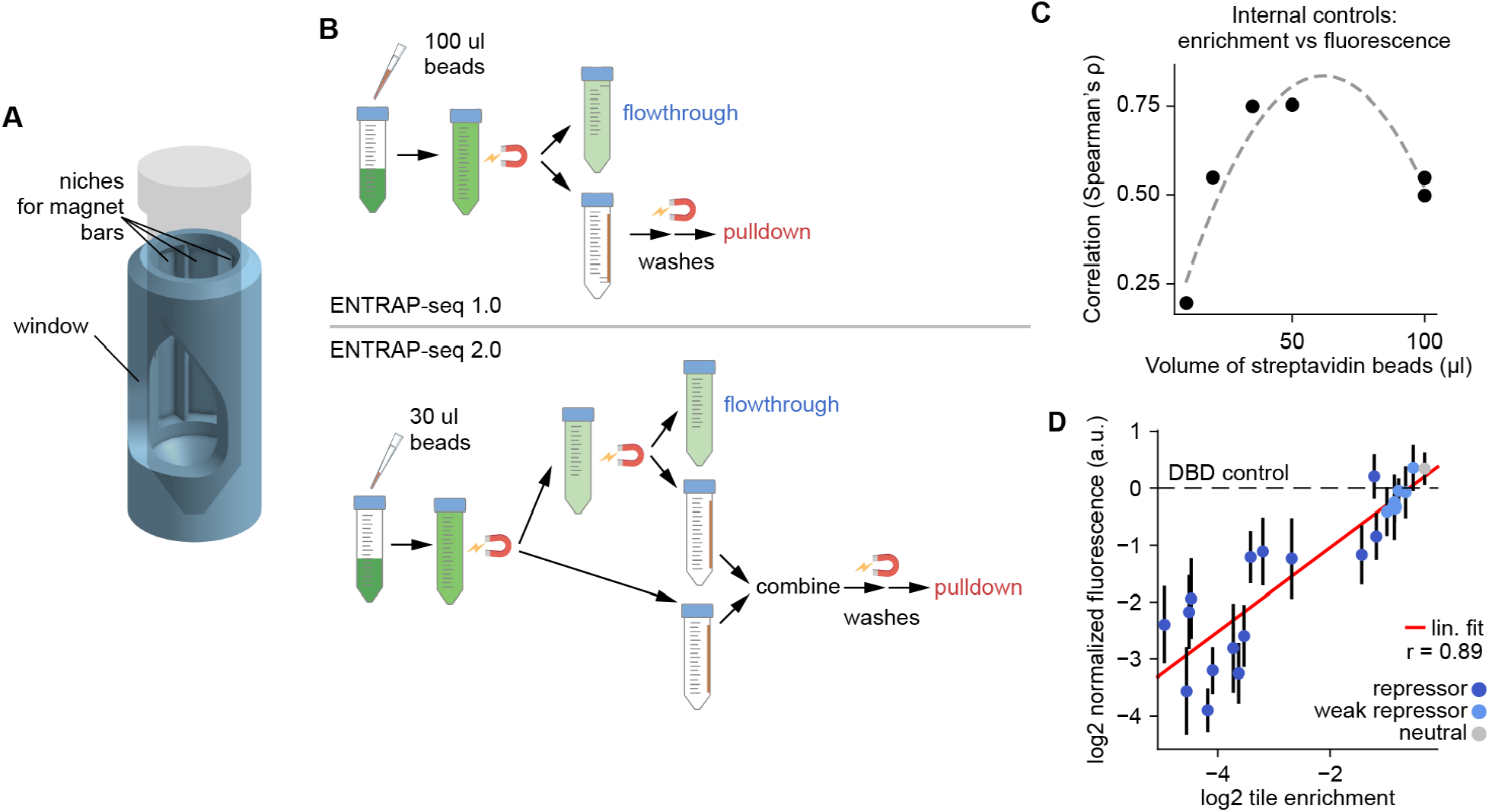
Optimization of ENTRAP-seq to simultaneously resolve repressors and activators. **(A)** Three-dimensional rendering of a custom-designed 3D-printed magnetic rack used for pulling down nuclei in a standardized fashion. **(B)** Schematic comparison of the original ENTRAP-seq protocol (ENTRAP-seq 1.0: top) and the new optimized protocol (ENTRAP-seq 2.0: bottom). In version 2.0, 30 µL of magnetic beads are added instead of 100 in order to decrease nonspecific binding. In addition, the new optimized protocol subjects the flowthrough of the first pulldown step to a second pulldown in order to remove remnant activators. Then, all pulldowns are combined and washed together. **(C)** Effect of the volume of magnetic streptavidin beads on assay accuracy. Library ATF6 was purified using 10, 20, 30, 50, and 100 µL of beads. Shown is the correlation between the activity of 50 internal controls measured using ENTRAP-seq versus a fluorescence reporter assay. The minimum bead volume that maximized the correlation was determined to be 30 µL and was used throughout this study. **(D)** Optimized ENTRAP-seq can quantitatively resolve the activity of newly identified repression domains. We selected tiles spanning a broad range of negative enrichment values and measured activity individually using a fluorescence reporter assay. Shown is the mean ± SD fluorescence (y-axis) as a function of tile enrichment (x-axis). N = 24 (4 plants with 6 leaf discs per plant).

**Supplementary Figure 3.**
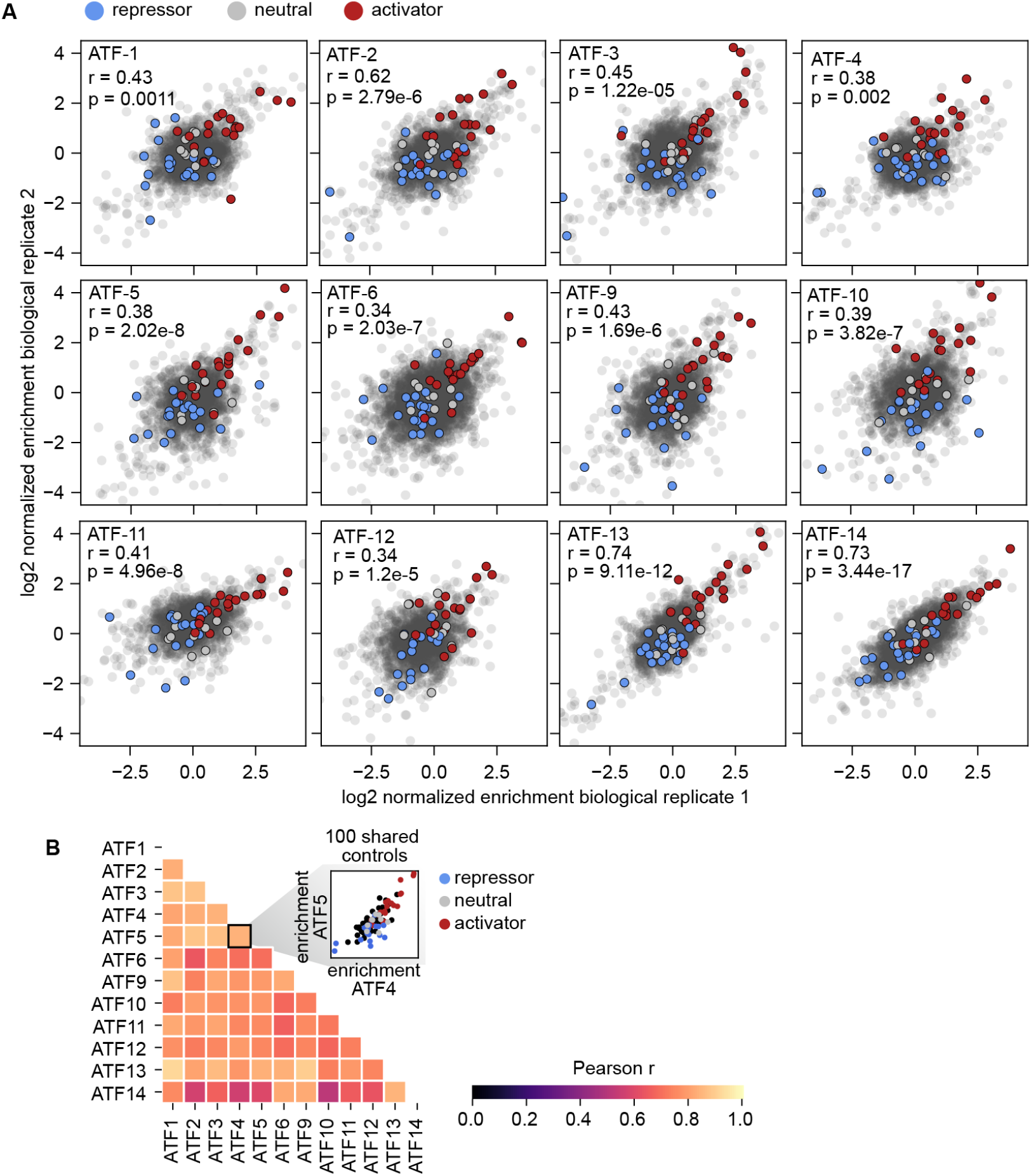
Biological reproducibility of ENTRAP-seq assays. **(A)** Scatter plots showing Z-score normalized tile enrichment (Methods) in two biological replicates of each ATF library. The Pearson correlation (r) and the p-value of the correlation coefficient (p) are indicated for each library. Shared internal controls are highlighted according to their activity class as repressors (blue), neutral (gray), and activators (red). **(B)** Pairwise Pearson correlations of mean enrichment values for the 100 shared internal controls across libraries. Inset shows an example comparison between libraries ATF4 and ATF5.

**Supplementary Figure 4.**
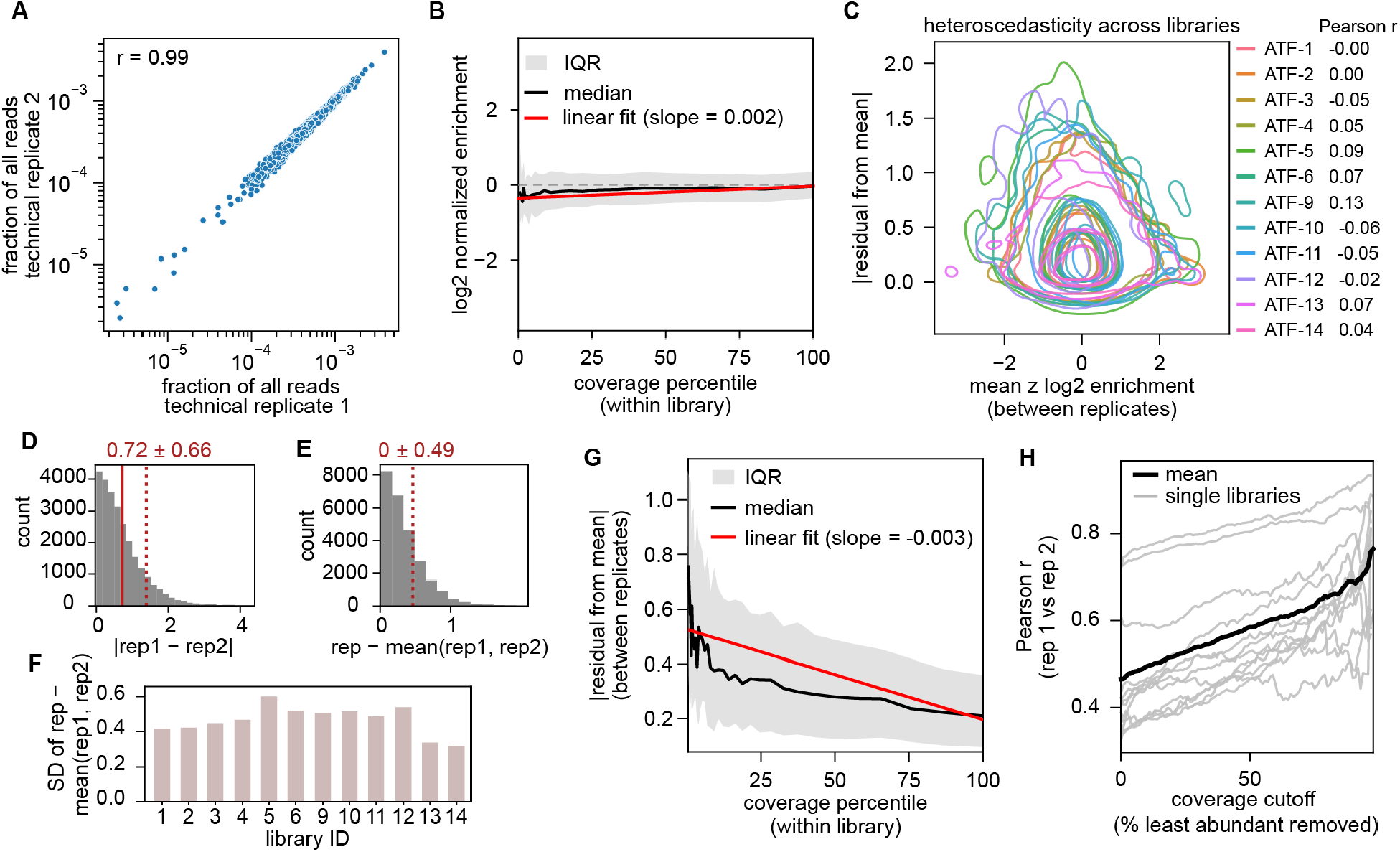
Assay reproducibility and sources of uncertainty. **(A)** Scatter plot showing the fraction of reads mapping to each tile in two technical replicates of library ATF4. Each replicate was prepared into a sequencing library independently and sequenced separately. Shown is the Pearson correlation between technical replicates (r=0.99). Most variability arises upstream of sequencing, potentially during agroinfiltration, nuclear isolation, and sorting. **(B)** Tile enrichment has an extremely weak correlation with tile representation in the library. Tile activity measured by ENTRAP-seq (y-axis) as a function of tile abundance in the library (x-axis, expressed as coverage quantile), for all tiles across all libraries. The solid black line shows the median across coverage quantile bins, the solid red line shows a linear fit to the data. There is a Δ=0.2 log2 enrichment units between the rarest and most abundant 1% of tiles. **(C)** Measurement error does not depend on tile enrichment. Density contour plot showing the relationship between mean tile enrichment between biological replicates (x-axis) and its error, quantified as the absolute residual from the mean (y-axis). The correlation between these two metrics for each library (i.e. heteroscedasticity) is shown as Pearson correlation coefficients. **(D)** Assessment of the measurement error. Shown is the distribution of absolute replicate differences across all libraries. **(E)** Assessment of the measurement error. Shown is the distribution of residuals to the mean of two biological replicates. **(F)** Estimation of the magnitude of the error. We start from the assumption that across all tiles the average enrichment μ between replicate 1 and replicate 2 is the best available estimate of true tile enrichment and that both replicates have the same standard deviation *σ* from this truth (i.e equal noise). Treating μ as the best estimate of truth for each tile, we ask how far does each replicate tend to sit from this mean. To answer this question we calculate the standard deviation of the residuals across all tiles, which captures the characteristic size of the noise. **(G)** Relationship between tile measurement error expressed as the residual from the biological replicate mean (y-axis) and tile coverage in the library (x-axis). Rare tiles have an appreciably higher error than abundant tiles. **(H)** The Pearson correlation between biological replicates of each library is plotted against the coverage cutoff used to filter out rare tiles. Removing rare tiles increases the reproducibility in a semi-linear fashion.

**Supplementary Figure 5.**
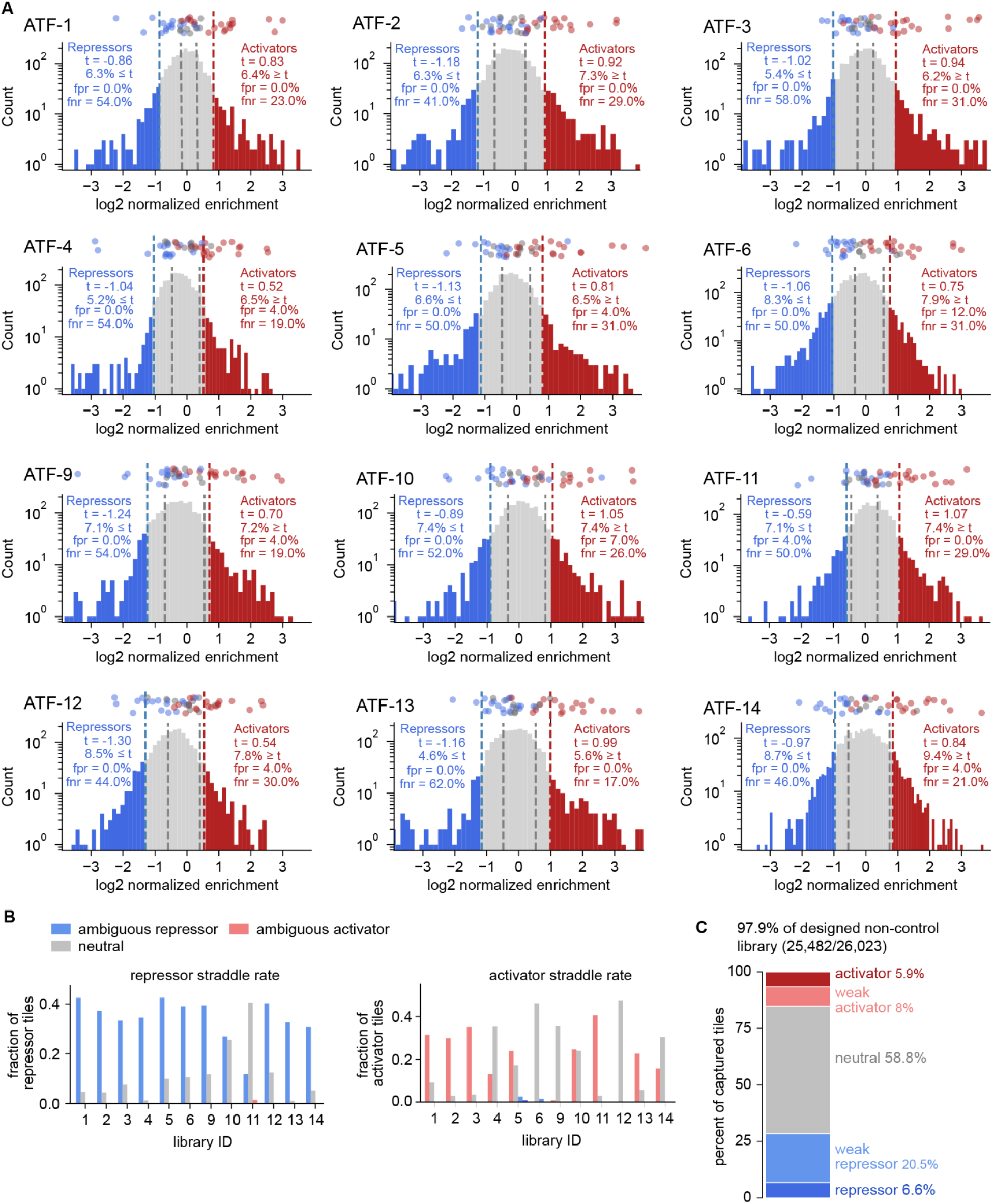
Qualitative classification of tiles into activity categories using internal controls. **(A)** Histograms of mean normalized tile enrichment obtained by averaging biological replicates and using the internal controls for z-score normalization. Shown on top are the values for 50 individual internal controls independently measured one by one using a fluorescence reporter assay (see Fig. S1F). Blue dashed lines correspond to the threshold for classifying tiles as repressors at -1.25 SD from the mean of all tiles in that library after control-based normalization. Red dashed lines correspond to the threshold for classifying tiles as activators at +1.25 SD from the normalized library mean. Gray dashed lines indicate ± 1SD from the mean of the neutral internal control tiles. In each library the legend shows t, the enrichment value at the 1.25 SD threshold, the percentage of tiles that lie beyond t, the false positive rate (fpr) and the false negative rate (fnr). The false positive and false negative rates are estimated using the validated internal controls. **(B)** Straddle rate for repressors (left) and activators (right), defined as the fraction of tiles in which one of the replicates had an absolute enrichment value lower than the classification threshold. The activity category of straddler tiles is shown as different colors. Note that by definition the absolute average enrichment of straddler tiles was higher than the threshold. **(C)** Stacked bar plot showing the percentage of captured non-control tiles belonging to each functional category across all libraries combined.

**Supplementary Figure 6.**
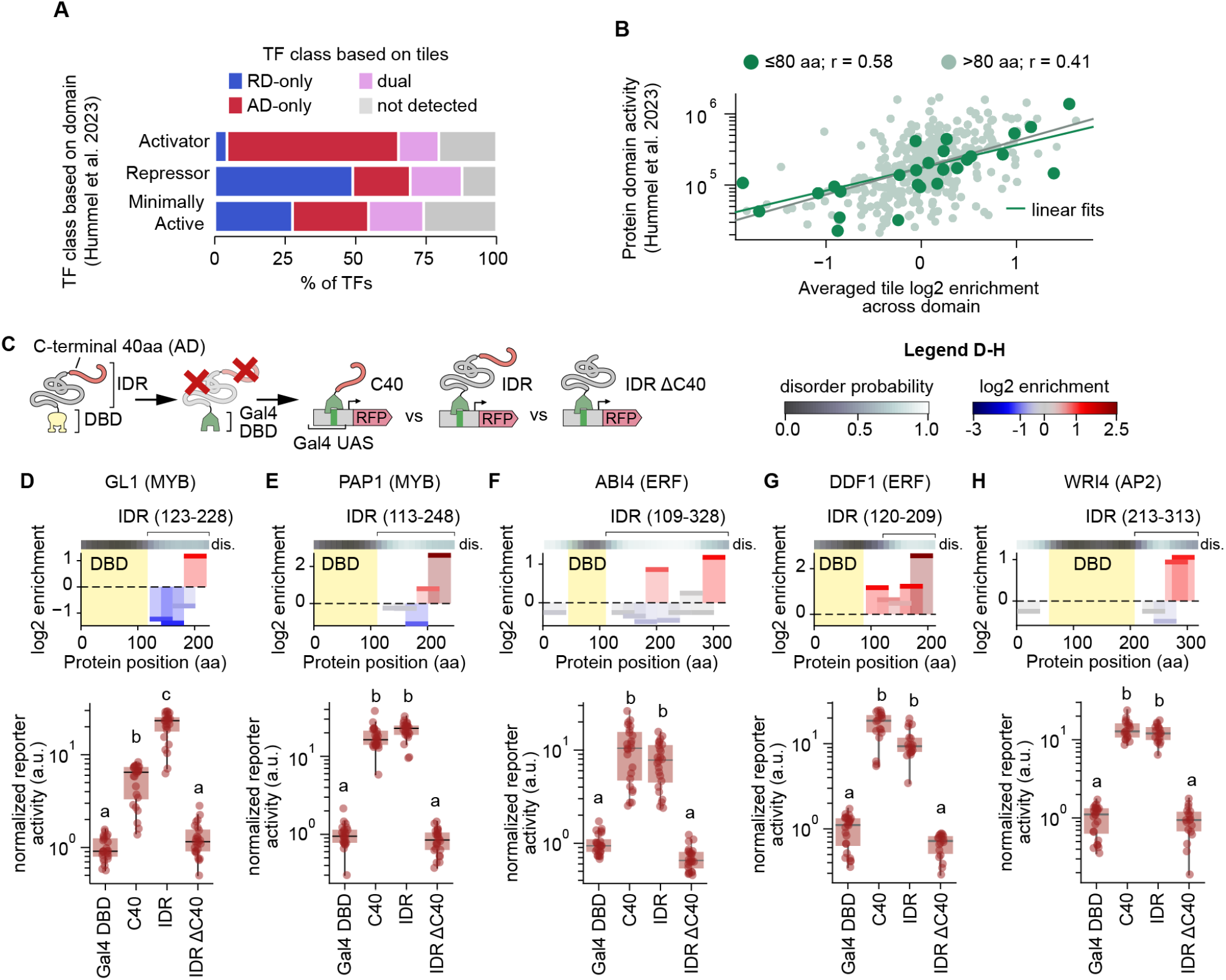
Quantitative validation of newly identified ADs and RDs. **(A)** Comparison of regulatory classifications obtained from non-tiled, non-DBD TF regions and 40-aa tiles. Hummel et al. previously tested 403 putative transcriptional effector domains, defined by excluding conserved DBDs and selecting the longest remaining contiguous non-DBD region from each TF. These regions were classified by the authors as activators, repressors, or minimally active (*28*). The same TF regions were reclassified here based on their constituent tiles using the criteria in Figure 1. The stacked bars show what percentage of full-length domain categories fall into each tile-based category. **(B)** Y-axis: regulatory activity of TF regulatory regions as measured individually using a fluorescence reporter assay (from (*28*) as in (A)). X-axis: activity of the regulatory regions measured as the average log2 enrichment of its constituent tiles. **(C)** Schematic of the experiments used to validate newly identified activation domains. To test necessity, C-terminal IDRs with their C-terminal 40-aa ADs intact or removed were fused to Gal4-DBD to drive a fluorescent reporter. To test for sufficiency, the C terminal 40-aa were tested by themselves. **(D)-(H)** Top: tile enrichment along the length of the TFs tested. The gray colorbar shows the predicted protein disorder according to the Metapredict software (*74*). The sequence of the IDRs tested as Gal4-DBD fusions is indicated. Bottom: Fluorescence reporter activity of IDR truncations. In (D)-(H) boxplots show the median ± IQR, whiskers correspond to minimum and maximum values excluding outliers, N = 24 (4 plants with 6 leaf discs per plant), letters on top of boxplots indicate statistically different groups (P < 0.05 based on a Tukey post hoc honestly significant difference test).

**Supplementary Figure 7.**
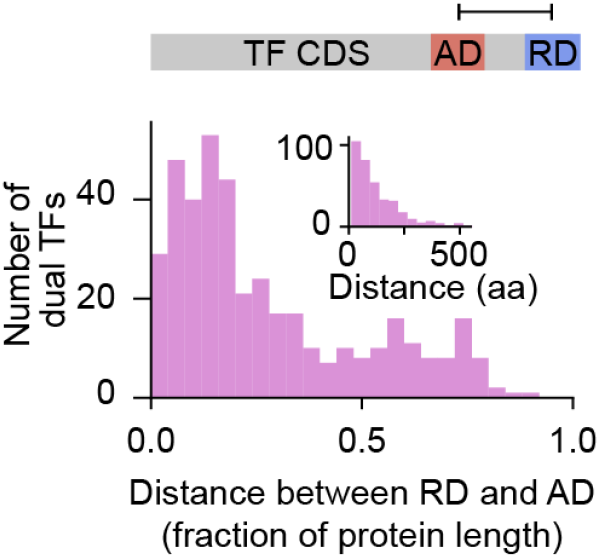
Separation between activation and repression domains in bifunctional transcription factors. Histogram showing the distance between tile centers of the closest activator and repressor tiles from dual TFs. Distance is expressed as the fraction of the length of each protein. The inset shows the absolute distance in amino acids.

**Supplementary Figure 8.**
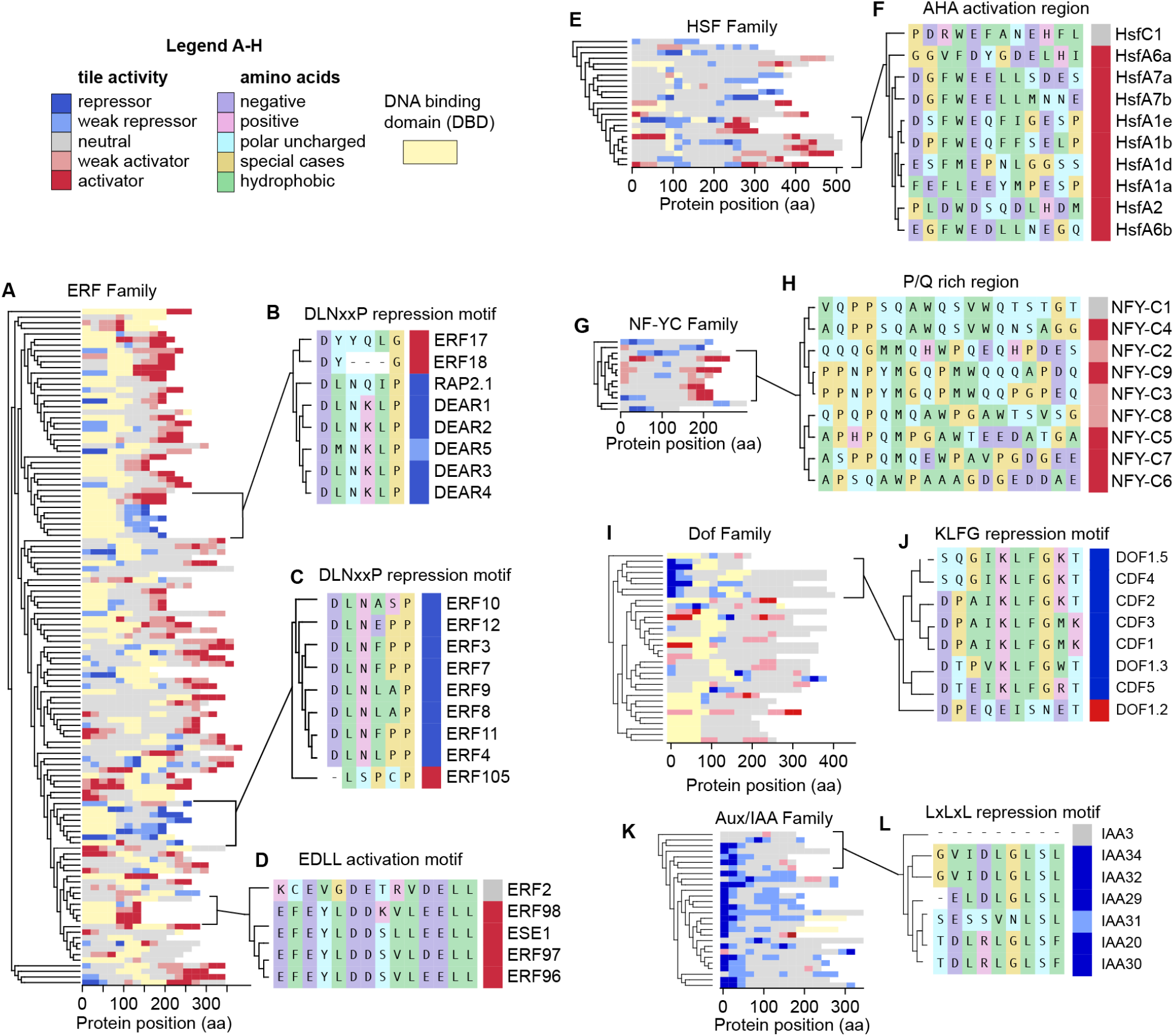
The Arabidopsis regulatory domain map recapitulates known repressor and activator TF clades and their domains. **(A)** Sequence-similarity dendrogram of the ERF family. The transcriptional activity of tiles along the length of each protein is shown. Tiles overlapping DBDs, which were not measured, are shown in yellow. **(B)** Multiple sequence alignment of the region containing the DLNxxP EAR motif among DEAR proteins and 2 closely related TFs. The transcriptional category of the corresponding tiles is shown in color-coded squares to the right of each sequence. The tiles carrying mutations in the motif are not repressors. **(C)** Same as (B) for a different clade of ERF repressors that appears to have independently evolved the DLNxxP repression motif. A closely related ERF protein, ERF105, has mutations in the motif and the corresponding tile is not a repressor. **(D)** Sequence alignment for TFs containing the EDLL activation motif and a closely related ERF TF where the motif is mutated and activation is absent from the corresponding tile. **(E)** Sequence-similarity dendrogram of the HSF family. **(F)** The core AHA activation motif among class A HSF members and a closely related HSF, HsfC1. The AHA motif has poor sequence conservation and it is not possible to connect specific mutations to the loss of activity in HsfC1. **(G)** Sequence-similarity dendrogram of TFs belonging to the NF-YC family. **(H)** Gapless alignment of the C-terminal PQ-rich region of a subset of NF-YC proteins. Like the HSF AHA domain, this region has poor linear sequence conservation and it is not immediately obvious why activation is lost in the NFY-C1 tile. **(I)** Sequence-similarity dendrogram of the Dof TF family. **(J)** Gapless alignment of the repression domain region in DOF TFs containing the KLFG repression domain. The most closely related DOF TF without an RD, DOF1.2, carries mutations in the conserved motif. **(K)** Sequence-similarity dendrogram of the Aux/IAA family. **(L)** Detail from the N-terminal RD in Aux/IAA proteins containing the LxLxL repression motif. This region is missing from IAA3 and the corresponding tile is transcriptionally neutral. IAA31 carries conservative substitutions in the motif and its corresponding tile is a weak repressor.

**Supplementary Figure 9.**
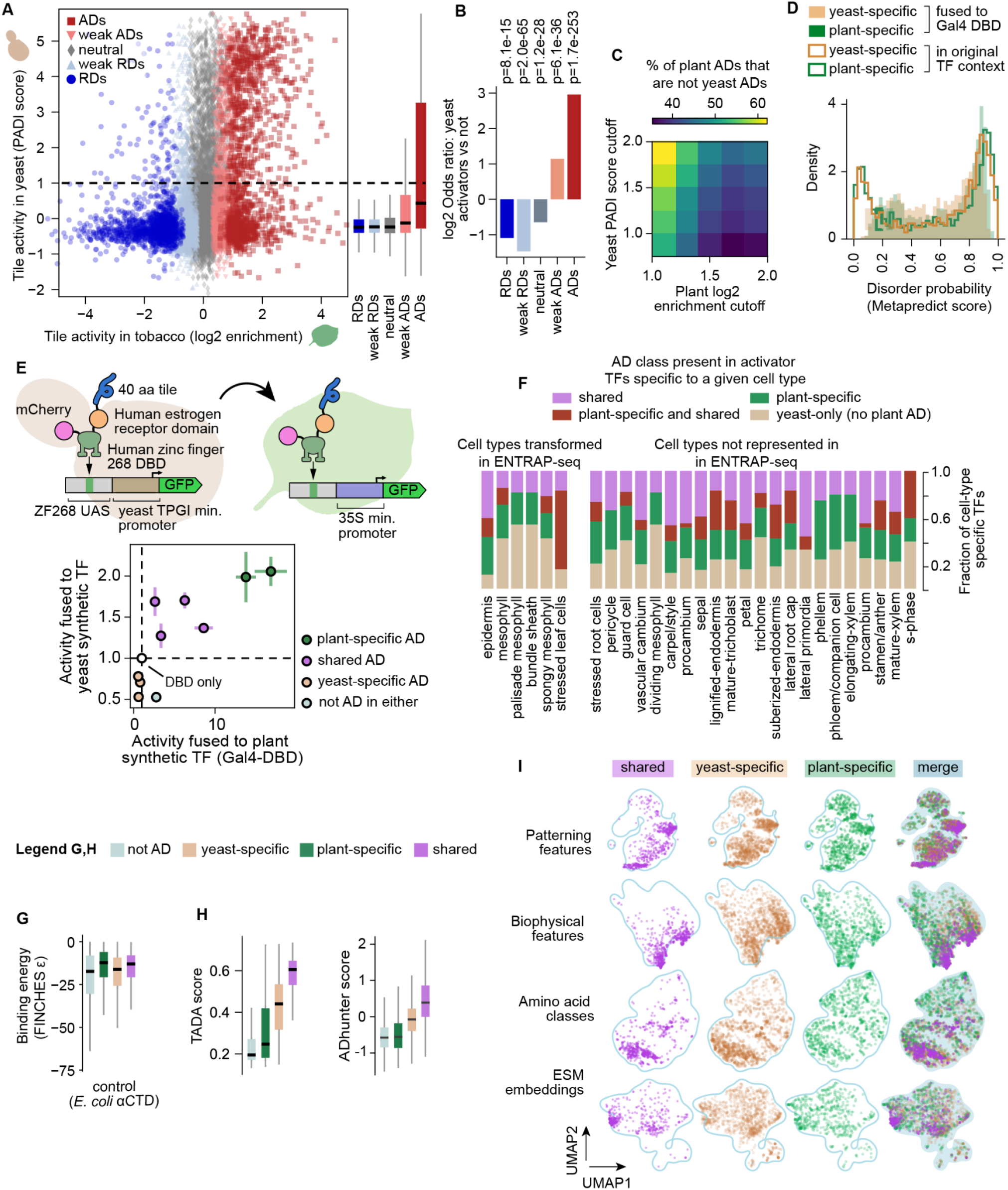
Partial overlap of activation domains between yeast and tobacco. **(A)** Scatter plot showing the transcriptional activity of Arabidopsis TF tiles in yeast (y-axis, PADI scores) and tobacco (x-axis, ENTRAP-seq enrichment). Tiles are color-coded according to their activity category in tobacco. Right: boxplots showing the PADI scores for each tobacco-based activity category. **(B)** Plant ADs are enriched in yeast ADs compared to non-ADs and the opposite is true for plant repressors. Bars show the odds ratio of each tile category versus the rest in yeast activators (PADI ≥ 1) versus yeast non-activators. P values on top correspond to a two-sided Fisher’s exact test. **(C)** A large fraction of ADs are plant-specific regardless of activation cutoffs. Heatmap showing what fraction of tiles classified as activators in tobacco are not activators in yeast as a function of the thresholds used to define activators in each experimental system. **(D)** Tile disorder does not explain differences in activity between tobacco and yeast. Density histogram of predicted protein disorder in yeast-specific and plant-specific ADs. Tile disorder was calculated using Metapredict on the tile as a Gal4-DBD fusion (filled histograms) or in their native TF context (contour histograms). **(E)** The synthetic TF used to recruit tiles to the reporter does not explain differences in activity between plants and yeast. The yeast synthetic TF was used to recruit tiles in a tobacco fluorescence reporter assay, the same tiles were also tested using the Gal4-DBD scaffold. The scatterplot shows the mean ± SD in activity using each synthetic TF normalized to either DBD alone. N = 24 (4 plants with 6 leaf discs per plant). **(F)** The fraction of yeast-only TFs is similar between cell types represented and unrepresented in ENTRAP-seq. TFs specific to a given cell type as defined by Baumgart et al. 2025 were classified according to the class of their ADs (*83*). TFs labeled as yeast-specific do not carry detected plant ADs and are similarly present in ENTRAP-seq cell types as other cell types. **(G)** Negative control for FINCHES modeling (Fig. 2). Boxplots showing the predicted binding energy between tiles of each AD class and the C-terminal domain of E. coli RNA Polymerase. **(H)** Activation scores for the ATF library tiles according to the TADA and ADHunter predictive models. **(I)** UMAPs of tiles belonging to the three AD categories projected onto different sets of sequence features. Outlines encompass 99% of all tiles. Box plots in (A), (G), and (H) show median ± IQR. Whiskers are minimum and maximum values excluding outliers.

**Supplementary Figure 10.**
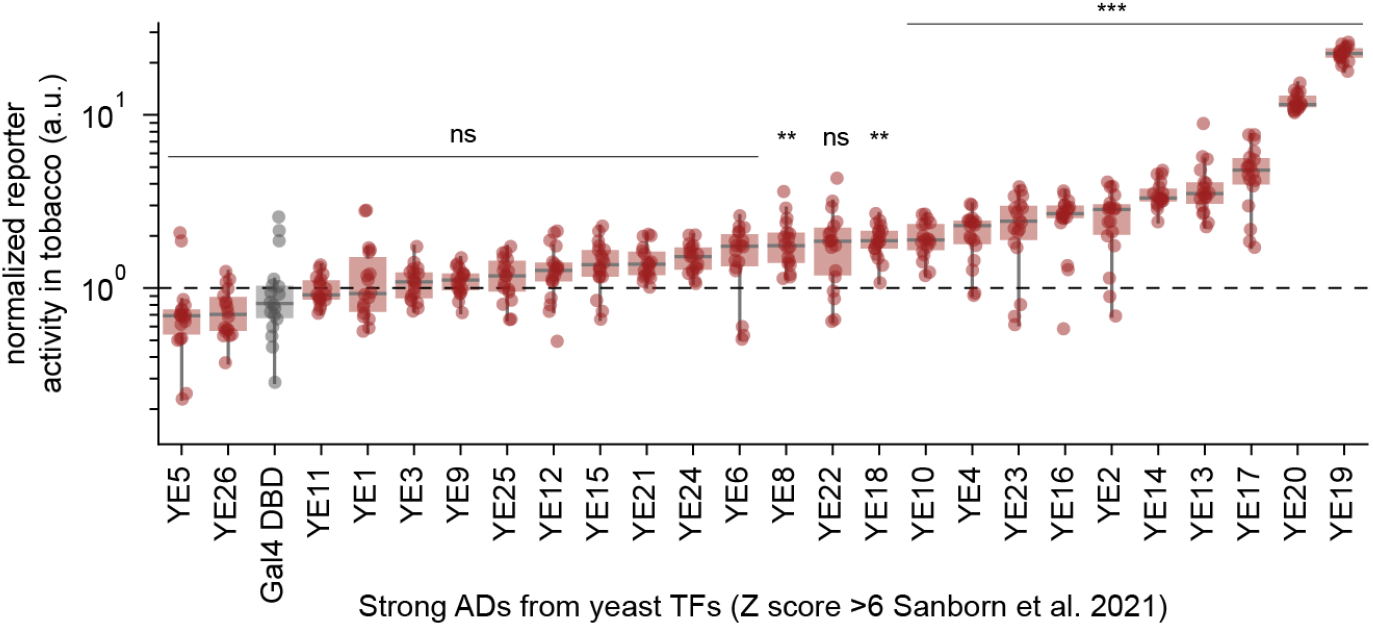
Many strong ADs from yeast TFs fail to activate in plants. A random sample of very strong 53-aa ADs from yeast TFs (Z score >6) were fused to the Gal4 DBD to drive a fluorescence reporter assay in tobacco leaves. Box plots show the median ± IQR, whiskers correspond to minimum and maximum values excluding outliers, N = 24 (4 plants with 6 leaf discs per plant). A Bonferroni-corrected two-sided Welch’s t-test against the Gal4-DBD control was used to classify fragments as ADs, *** p<0.001, ** p< 0.01, * p<0.05, ns not significantly different from Gal4-DBD.

**Supplementary Figure 11.**
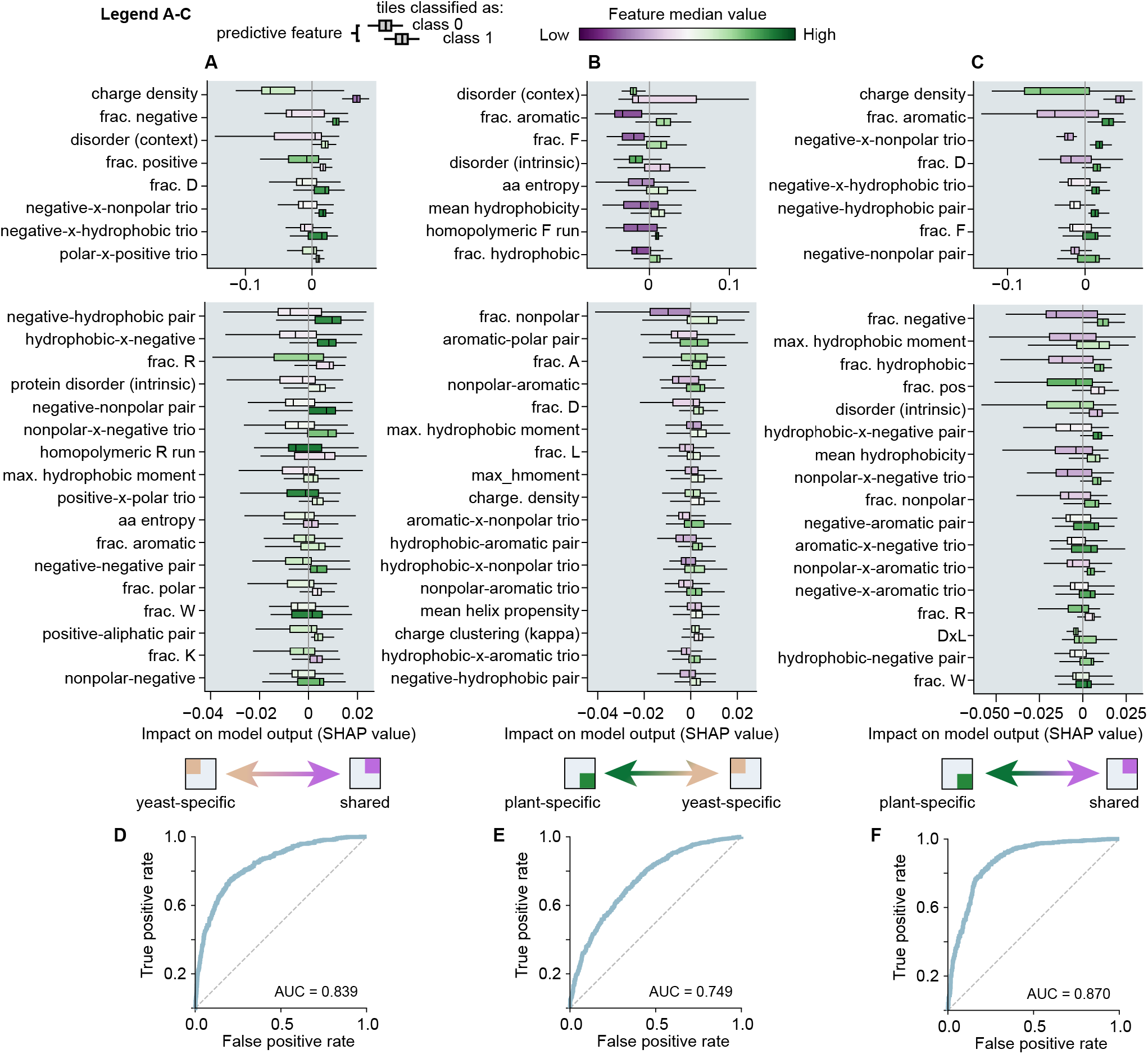
Sequence features driving the classification between activation domain classes in random forest classifiers. **(A-C)** Random forests were trained on sequence features to separate classes of ADs. Top: SHAP summary plot showing the contribution of the top 8 most predictive features to the random forest classifier separating two classes of ADs. Each pair of box plots corresponds to one feature. The top boxplot shows values for tiles classified as class 0 (i.e yeast-specific in (A)) and the bottom box plot shows SHAP values for tiles classified as class 1 (i.e shared ADs in (A)). The hue of the box plots corresponds to the median value of that feature in each class. Bottom: SHAP summary plot for the 3rd-25th most predictive features. Box plots show the median ± IQR, whiskers correspond to minimum and maximum values excluding outliers. **(D-F)** Receiver Operating Characteristic (ROC) curves for each classification. The area under the curve (AUC) is indicated. Box plots show the median ± IQR, whiskers correspond to minimum and maximum values excluding outliers.

**Supplementary Figure 12.**
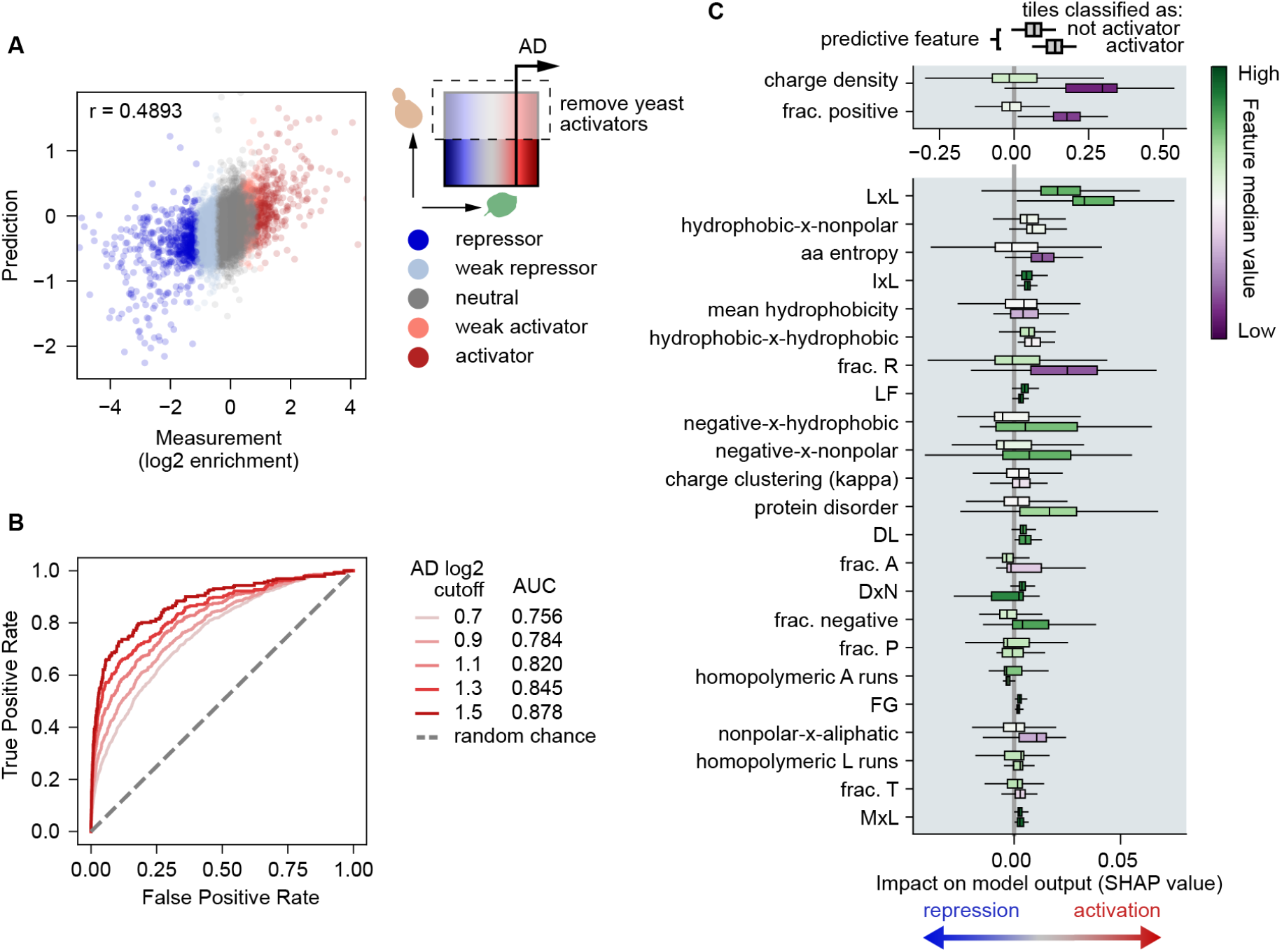
Predictive model for the plant-specific axis of activation. **(A-C)** XGBoost regressor ensembles were trained on tile features (Table S2) to predict ENTRAP-seq log2 enrichment values. Tiles classified as activators in yeast (PADI ≥1) were excluded (*4*). Data were split 80:20 into 5 cross-validation folds. **(A)** Predicted (y-axis) and observed (x-axis) ENTRAP-seq log2 enrichment values of out-of-fold test data. Points are colored according to the experimental classification of each tile. **(B)** Classification performance was evaluated using Receiver Operating Characteristic Area Under the Curve (ROC/AUC) on all out-of-fold predictions. To this end, the ground truth for tiles belonging to the activator class was defined based on increasing enrichment cutoffs to assess classification performance as a function of activator strength. Listed are the AUC values for increasingly stringent definitions of the activator class. **(C)** Summary SHAP plot showing the contribution of the 25 most predictive features to the out-of-fold predictions made across all regressors used in cross-validation in (A) and (B). Each pair of box plots corresponds to one feature. The top boxplot shows values for tiles classified as class 0 (i.e not AD) and the bottom box plot shows SHAP values for tiles classified as class 1 (i.e AD). The hue of the box plots corresponds to the median value of that feature in each class. Box plots show the median ± IQR, whiskers correspond to minimum and maximum values excluding outliers.

**Supplementary Figure 13.**
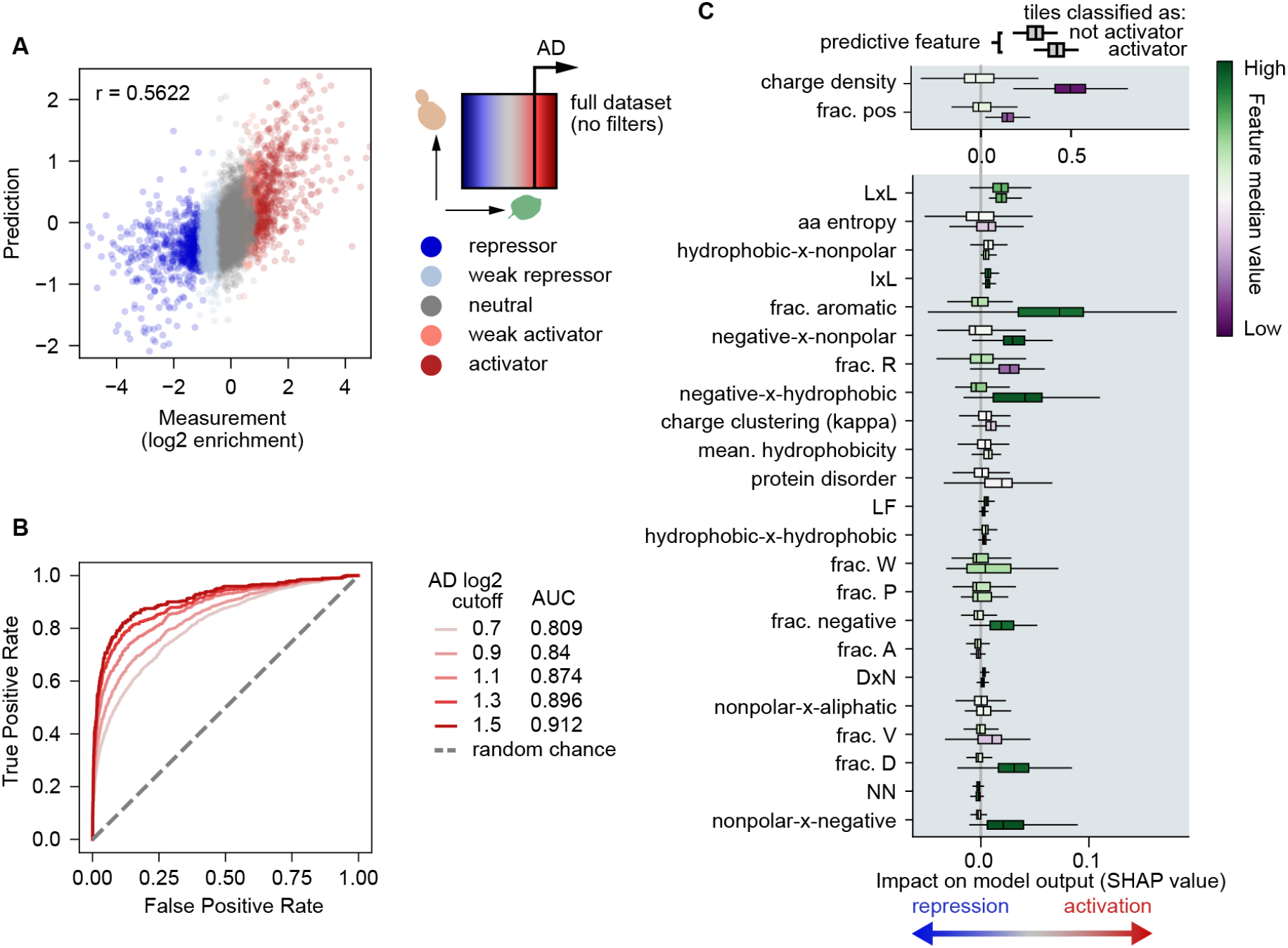
General predictive model of the complete dataset. **(A-C) (A-C)** XGBoost regressor ensembles were trained on tile features (Table S2) to predict ENTRAP-seq log2 enrichment values. Unlike Fig. S12, tiles classified as activators in yeast (PADI ≥1) were also included (*4*). Data were split 80:20 into 5 cross-validation folds. **(A)** Predicted (y-axis) and observed (x-axis) ENTRAP-seq log2 enrichment values of out-of-fold test data. Points are colored according to the experimental classification of each tile. **(B)** Classification performance was evaluated using Receiver Operating Characteristic Area Under the Curve (ROC/AUC) on all out-of-fold predictions. To this end, the ground truth for tiles belonging to the activator class was defined based on increasing enrichment cutoffs to assess classification performance as a function of activator strength. Listed are the AUC values for increasingly stringent definitions of the activator class. **(C)** Summary SHAP plot showing the contribution of the 25 most predictive features to the out-of-fold predictions made across all regressors used in cross-validation in (A) and (B). Each pair of box plots corresponds to one feature. The top boxplot shows values for tiles classified as class 0 (i.e not AD) and the bottom box plot shows SHAP values for tiles classified as class 1 (i.e AD). The hue of the box plots corresponds to the median value of that feature in each class. Box plots show the median ± IQR, whiskers correspond to minimum and maximum values excluding outliers.

**Supplementary Figure 14.**
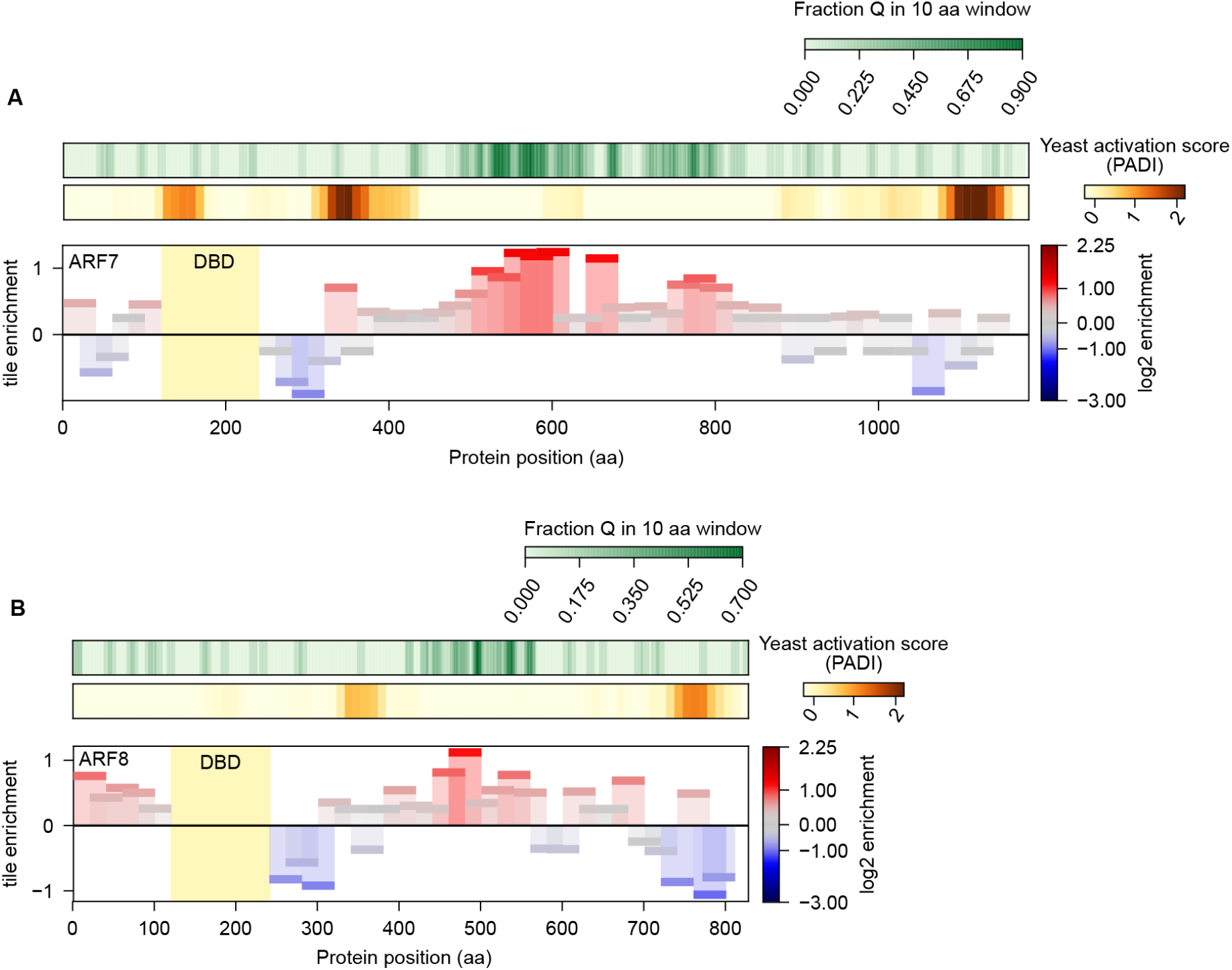
Identification of Glutamine-rich activation domains in the middle region of class A ARF activators. **(A)** Arabidopsis ARF7 protein showing top: percent glutamine in a 10 aa moving window, middle: transcriptional activity of 40-aa tiles in yeast, bottom: transcriptional activity of 40-aa tiles in tobacco measured by ENTRAP-seq. **(B)** Same as (A) for ARF8.

**Supplementary Figure 15.**
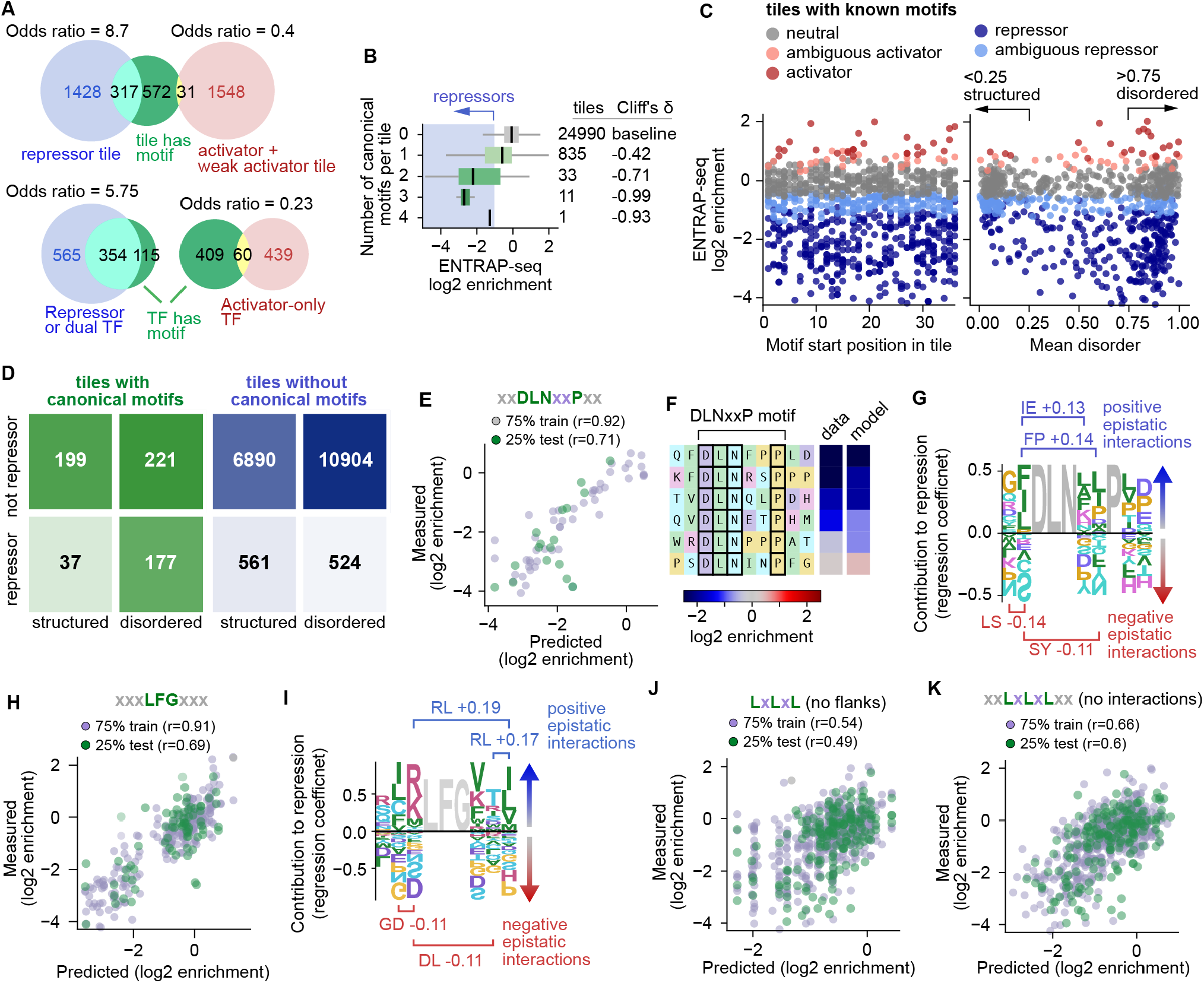
Systematic dissection of canonical repression motifs: additional details. **(A)** Strong but incomplete linkage between motifs and tile activity. Top: Venn diagrams show the overlap between tiles having a canonical motif and them being repressors or activators. Bottom: Venn diagrams show the overlap between TFs being classified as repressor or dual –i.e. having an RD– and the TF having a canonical repression motif anywhere in its corresponding tiles. The same is shown for activator TFs. **(B)** Boxplots of ENTRAP-seq enrichment for tiles containing 0-4 canonical repression motifs. **(C)** Tile activity (ENTRAP-seq enrichment, y-axis) for motif-containing tiles as a function of the position of the motif along the tile (left), or the predicted tile structural disorder (right). **(D)** Truth tables of activity (rows, repressor or not repressor) versus disorder (columns, Metapredict predicted disorder < 0.25 or Metapredict predicted disorder > 0.75). **(E)** Measurements (y-axis) versus predictions (x-axis) for a model of the EAR motif DLNxxP with two flanks. **(F)** Examples of motifs from Arabidopsis tiles displaying measured and predicted differences in repression strength. Consensus positions are outlined. **(G)** Repression weights logo for DLNxxP with two flanks. The height of each amino acid corresponds to the magnitude of its linear weight either contributing to repression (positive weights), or against it (negative weights). The two strongest pairwise interactions favoring or disfavoring repression are also shown. **(H)** Measured versus predicted enrichment of tiles containing the LFG tripeptide, part of the canonical R/KLFGV motif. **(I)** Repression weights logo from the LFG tripeptide model. **(J)** Flanking residues contribute to EAR motif strength prediction. Scatter plot shows ENTRAP-seq measurements (y-axis) versus model predictions (x-axis) for the LxLxL motif without flanks. **(K)** Pairwise interactions between free positions contribute to EAR motif strength prediction. Scatter plot shows measurements (y-axis) against predictions (x-axis) for the LxLxL EAR motif with 2 flanks but no pairwise interactions between free positions.

**Supplementary Figure 16.**
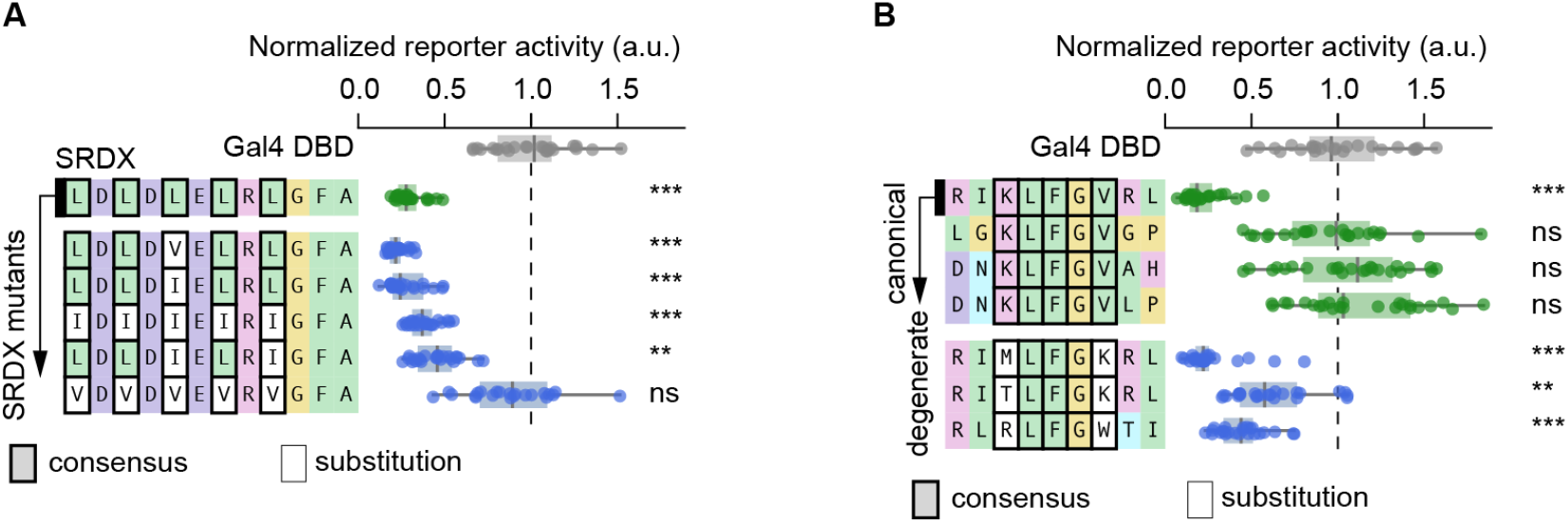
Sequence context can matter as much as consensus positions. **(A)** Fluorescence reporter assay of Gal4 DBD fusion peptides encoding variants of SRDX, a potent and widely known synthetic LxLxL EAR motif. On top, the original SRDX peptide in green. SRDX mutants carrying substitutions in consensus L are shown in blue at the bottom. **(B)** Fluorescence reporter assay of Gal4 DBD fusion peptides encoding variants of the canonical KLFGV motif ± 2 flanking residues. On top in green are canonical variants with a conserved KLFGV core but variant flanks. The strongest canonical variant was used as a template for degenerate variants carrying substitutions in consensus positions (bottom in blue). Box plots in (A) and (B) show the median ± IQR, whiskers correspond to minimum and maximum values excluding outliers. *** p<0.001, ** p< 0.01, * p<0.05, ns not significantly lower using a one-sided Welch’s t-test against Gal4-DBD control. N = 24 (4 plants with 6 leaf discs per plant).

**Supplementary Figure 17.**
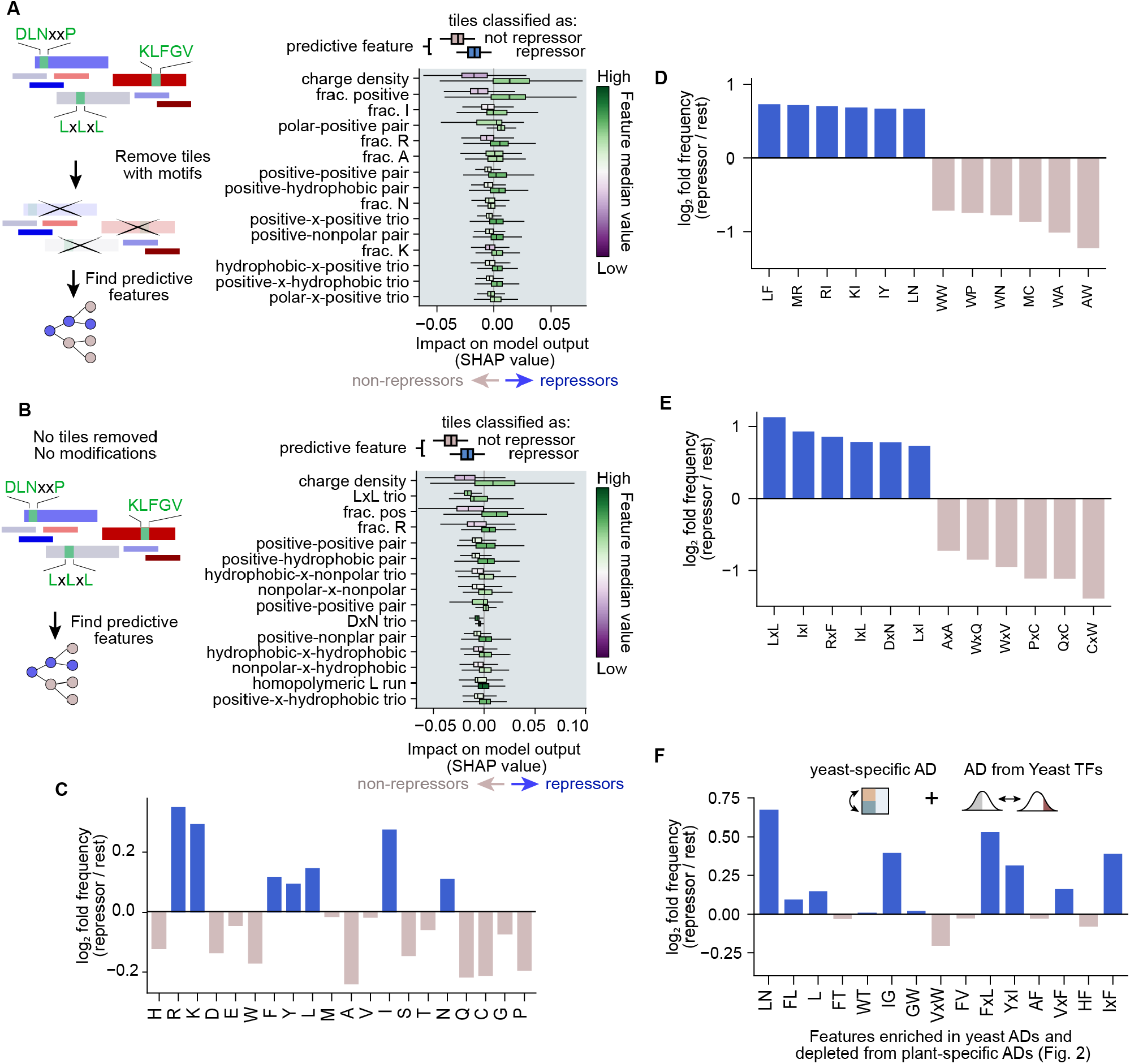
Protein features associated with repression domains: additional details. **(A)** Left: Tiles containing canonical motifs were removed from the dataset, independent of their activity. Biophysical and sequence features were used to train a random forest classifier to classify repressors from nonrepressors (neutral, weak activators, and activators). Right: Summary SHAP plot showing the contribution of the 15 most predictive features to the held-out test set. Each pair of box plots corresponds to one feature. The top boxplot shows values for tiles classified as not repressor and the bottom box plot shows SHAP values for tiles classified as repressor. The hue of the box plots corresponds to the median value of that feature in each class. **(B)** Same as (A) except no tiles were removed from the dataset. In (A) and (B) Box plots show the median ± IQR, whiskers correspond to minimum and maximum values excluding outliers. **(C)** Fold change in the frequency of individual amino acids in repressors compared to nonrepressors (neutral, weak activators, and activators). **(D)** Fold change in the frequency of dipeptides in repressors compared to nonrepressors. Only the top 6 most enriched and most depleted are shown. **(E)** Fold change in the frequency of gapped dipeptides in repressors compared to nonrepressors (x = any amino acid). Only the top 6 most enriched and most depleted are shown. **(F)** Listed on the x-axis are amino acids and dipeptides enriched in yeast activators and simultaneously depleted from plant-specific activators (Fig. 2). For each of these features, the y-axis shows the enrichment or depletion in repressors versus nonrepressors.

**Supplementary Figure 18.**
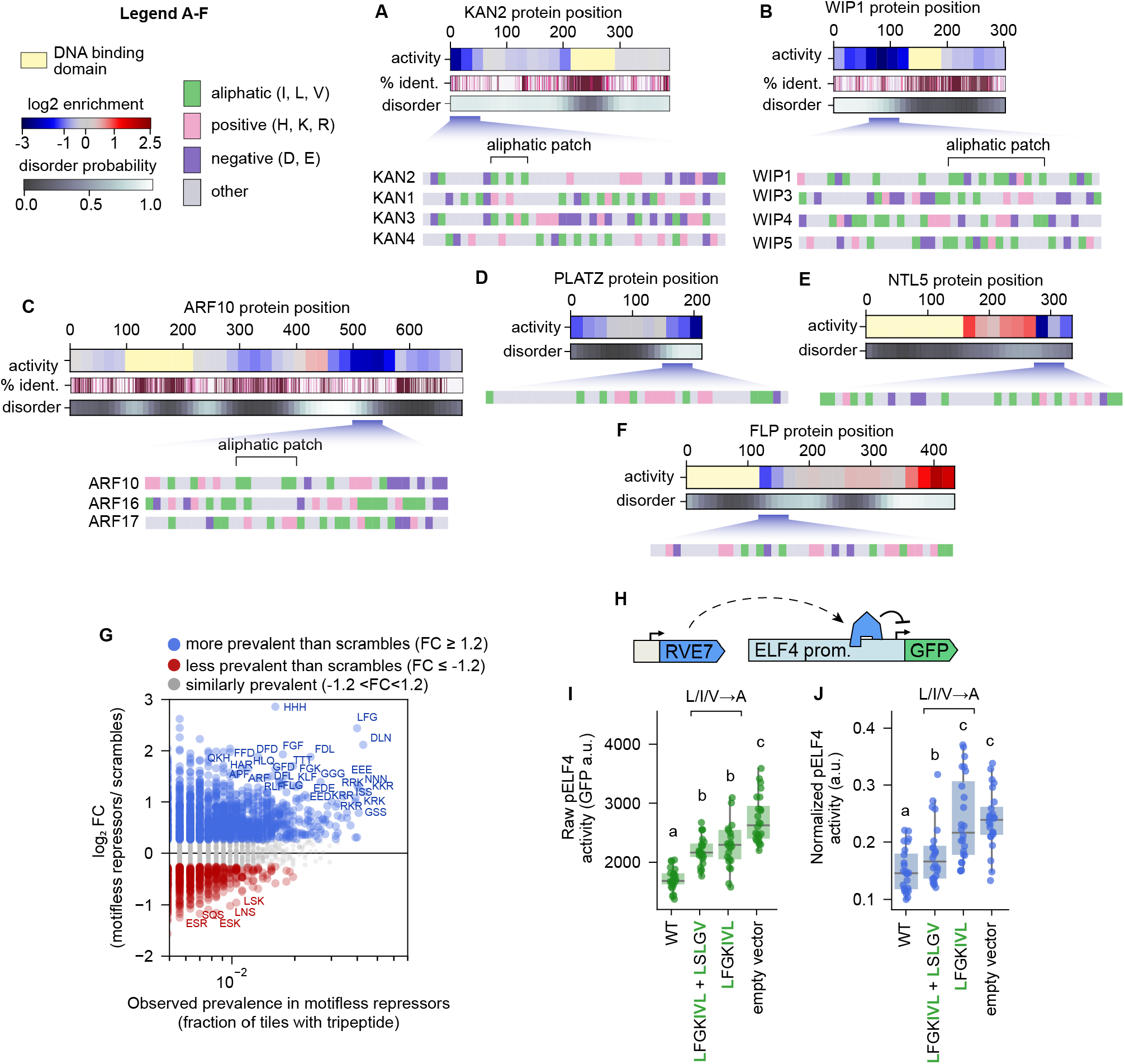
Sequence features of motifless repression domains. **(A)-(C)** Additional examples of protein families containing RDs with high prediction scores. These RDs are usually located in densely charged tiles with an overall positive charge and contain clusters of L, I, or V residues (aliphatic patches). **(D)-(F)** Additional examples of singleton proteins containing RDs with high prediction scores. These proteins do not have a homolog TF with a homologous motifless RD. **(G)** Abundance of tripeptides in motifless RDs with respect to 50 composition-preserving scrambles per tile. The x-axis shows the fraction of motifless RDs that contain a tripeptide, the y-axis shows the log2 fold change between that fraction and the fraction across the scrambled library. **(H)** Schematic of the RVE7 reporter assay. The full-length RVE7 protein was coexpressed with a GFP reporter driven by the Arabidopsis ELF4 promoter, an endogenous target repressed by RVE7. **(I)** Raw (not normalized) activity of the ELF4 promoter coexpressed with different RVE7 variants. Letters in green indicate residues in RVE7 that were mutated to A. **(J)** Same as (I) but normalizing the data to a constitutive BFP strain. In (I) and (J) boxplots show the median ± IQR, whiskers correspond to minimum and maximum values excluding outliers, N = 24 (4 plants with 6 leaf discs per plant), letters on top of boxplots indicate statistically different groups (P < 0.05 based on a Tukey post hoc honestly significant difference test).

**Supplementary Figure 19.**
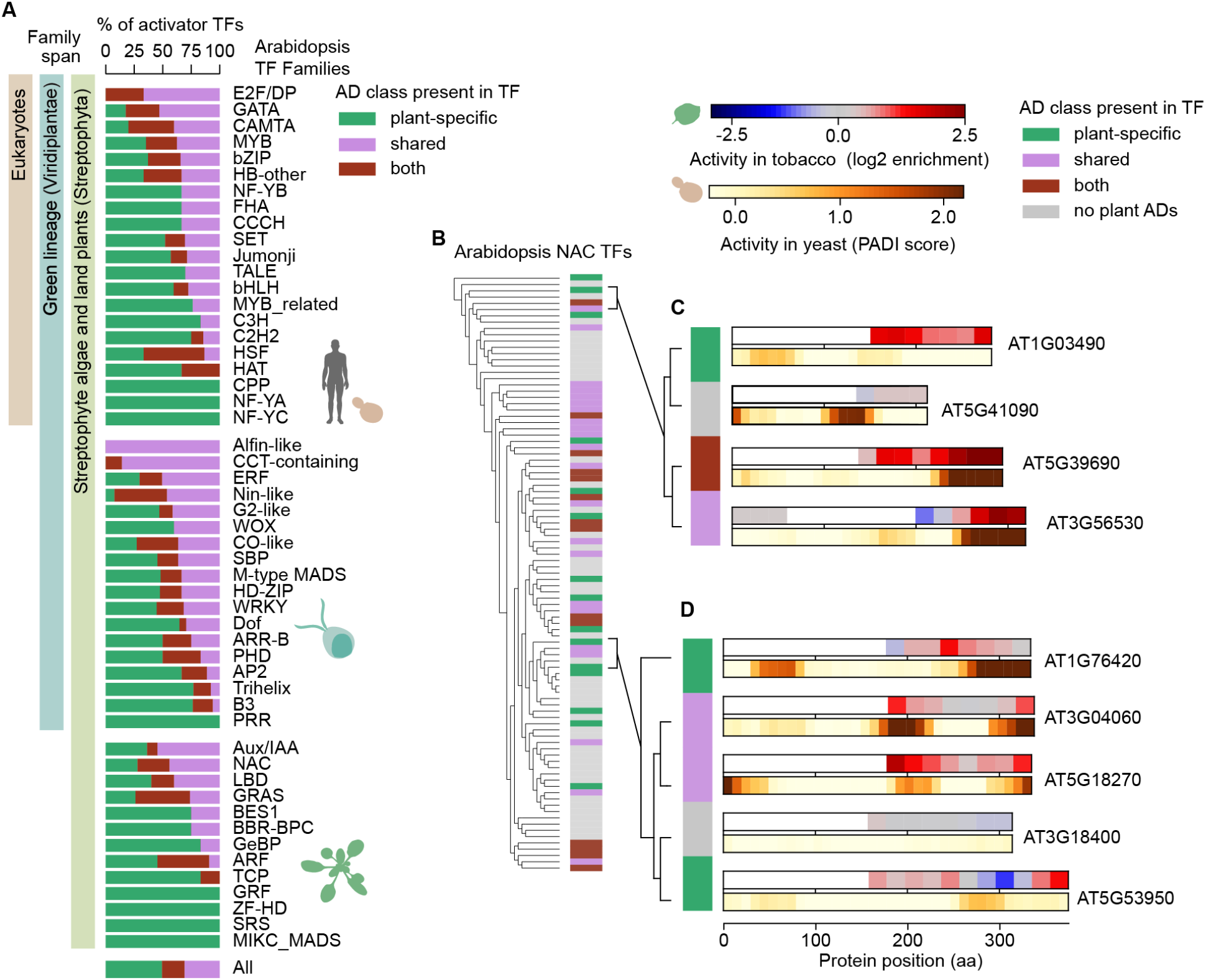
Evolutionary distribution of AD classes in Arabidopsis TFs. **(A)** TF families were manually grouped into three broad taxonomic-depth categories based on comparative surveys and family-specific phylogenetic studies (*69*, *84*, *85*): broadly eukaryotic TFs, TFs that are thought to have emerged in the green lineage (Viridiplantae) and TFs that are thought to have emerged in land plants and streptophyte algae (Streptophyta). The activator TFs of each family (that is, the TFs that have at least one tile that activates in tobacco) were classified into 3 groups according to whether their ADs also activate in yeast (pink) or are plant-specific (green). TFs labeled as both (brown) carry at least one plant-specific AD and at least one shared AD. **(B)** Sequence-similarity dendrogram of Arabidopsis NAC TFs classified using the same categories as in (A). TFs without ADs identified in tobacco appear in gray. **(C, D)** Subclades of NAC TFs showing tile activity along protein length in tobacco and yeast.

